# POMPOMS: Crosslinked biomolecular condensates as a versatile platform for multifunctional protein microparticles

**DOI:** 10.1101/2025.06.13.659498

**Authors:** Augene S. Park, Erika A. Ding, Benjamin S. Schuster

## Abstract

Protein-based microparticles are promising materials for applications such as biocatalysis and biomolecular capture, yet their fabrication by existing techniques remains challenging due to protein denaturation or lack of spatial control. Here, we present a method for synthesizing microscale protein-based materials by chemically crosslinking biomolecular condensates. Leveraging the liquid-liquid phase separation behavior of intrinsically disordered RGG domains, we sequestered RGG-tagged fusion proteins into droplets, then we solidified them into porous microparticles using the homobifunctional, amine-reactive crosslinker BS³. By modulating protein concentration and condensate coalescence, we controlled microparticle size from <1 to >40 µm. We then demonstrated three encodable functionalities: We used the SpyCatcher/SpyTag system to capture cargo proteins, we crosslinked core-shell condensates to generate microparticles with controlled spatial organization, and we immobilized a thermostable alcohol dehydrogenase with 31% retained enzymatic activity. These POMPOMS (protein-based, self-organized microparticles of multifunctional significance) represent a sustainable, tunable platform for versatile protein-based materials.

## INTRODUCTION

Intrinsically disordered proteins (IDPs) lack a defined tertiary structure and instead adopt an ensemble of interconverting conformations.^1–3^ The structural plasticity of IDPs allows many of them to undergo liquid-liquid phase separation (LLPS). Biomolecular LLPS is a thermodynamic phenomenon where proteins and other biomolecules demix into dense, liquid-like coacervates.^3,4^ IDP coacervation is governed by weak, multivalent interactions (e.g. hydrophobic, π-π, cation-π, and electrostatic interactions) that allow for reversible assembly.^5–7^ Within cells, phase-separated IDPs contribute to the formation of membraneless organelles known as biomolecular condensates, which can mediate spatial organization, regulate biochemical reactions, and dynamically sequester cellular components.^5–10^

Leveraging this dynamic behavior, researchers are now engineering IDPs into next-generation, protein-based materials.^11–16^ IDPs offer distinct advantages for this purpose compared to synthetic polymers. The monodispersity (i.e. uniform size and composition) of proteins ensures reproducibility.^17–20^ Recombinant protein expression allows for scalable synthesis of IDPs with biodegradability and reduced environmental persistence, addressing sustainability challenges associated with petrochemical-derived polymers.^14,21–23^ Phase-separating IDPs can be isolated by temperature-induced precipitation, bypassing costs and technical challenges associated with operating large-scale chromatography processes.^24–27^ Genetic engineering enables the precise customization of interaction motifs and phase behavior.^11,14,17,28^

While nanoscale (e.g. nanoparticles) and macroscale (e.g. hydrogels, fibers) protein-based materials are well-established, microscale protein-based materials – particularly microparticles – remain relatively unexplored despite their unique potential.^29^ Microparticles combine the high surface area-to-volume ratios inherent to the colloidal regime with the practical usability of macroscale materials. Much of the existing published literature on protein-based microparticles has so far focused on a limited number of applications such as the encapsulation of bioactive molecules for food production and drug delivery.^12,30–33^ However, protein-based microparticles can be potentially used in a wider array of applications like biomolecular capture and enzyme immobilization. Microparticles with interaction motifs could be used to purify products in biomanufacturing workflows or sequester molecules of interest for downstream analysis in fields such as diagnostics or pollution detection. Protein-based microparticles could also be engineered as immobilized enzyme systems, which are important for biocatalytic industrial processes, especially when the biocatalyst needs to be separated from the products at the conclusion of the reaction.

The dearth of research on protein-based microparticles may be due to constraints associated with current biopolymer material synthesis techniques. Proteins can naturally self-assemble to form nanoscale structures, but many have difficulty forming micrometer-sized materials with a well-defined morphology.^34^ Commonly used macroscale techniques such as casting or extrusion lack the fine spatial control required to create smaller, micrometer-sized materials. Existing microscale approaches such as spray drying, solvent extraction, 3D printing, and inkjet printing often use conditions (e.g. high temperature, organic solvents) that risk protein denaturation.^33,35–37^ The limitations of these techniques also restrict the types of proteins that can be used for microparticle synthesis.

To address challenges of fabricating protein-based microparticles and to expand the repertoire of proteins amenable for bioactive material synthesis, we developed a facile method for forming protein microparticles by chemically crosslinking biomolecular condensates. We fused intrinsically disordered RGG domains, derived from the *C. elegans* LAF-1 protein, to folded proteins to promote phase separation.^38,39^ The RGG domain, whose phase behavior we and others have extensively characterized, is a 168-residue domain rich in arginine-glycine-glycine motifs and drives phase separation via multivalent interactions under low-salt and low-temperature conditions.^39–41^ By inducing condensate formation with the RGG-tagged fusion proteins, we sequestered proteins in close enough proximity to be crosslinked to each other. Then we crosslinked the condensates using the homobifunctional, amine-reactive crosslinker bis(sulfosuccinimidyl) suberate (BS³) (Fig. 1A). To validate our approach, we crosslinked RGG-GFP-RGG biomolecular condensates and used confocal microscopy to observe the change in material properties from liquid to solid upon crosslinking. We quantified how varying the crosslinking conditions affords control over the size of crosslinked condensates. Then, we extended our strategy to demonstrate three functionalities of crosslinked biomolecular condensates: First, we crosslinked RGG-SpyCatcher-RGG condensates for molecular capture of SpyTagged cargo proteins. Second, we engineered hierarchical microparticle structure by using designed surfactant proteins that decorate the surface of crosslinked condensates. Third, we crosslinked alcohol dehydrogenase-RGG condensates as proof-of-concept for a novel method of enzyme immobilization. Overall, this work establishes crosslinked condensates as a versatile platform for functional, porous protein microspheres. We call this system POMPOMS – <u>p</u>rotein-based, self-<u>o</u>rganized <u>m</u>icro<u>p</u>articles <u>o</u>f <u>m</u>ultifunctional <u>s</u>ignificance. By bridging the gap between nanoscale precision and macroscale practicality, POMPOMS unlock applications in fields such as biocatalysis and biomolecular capture.

**Figure 1.**
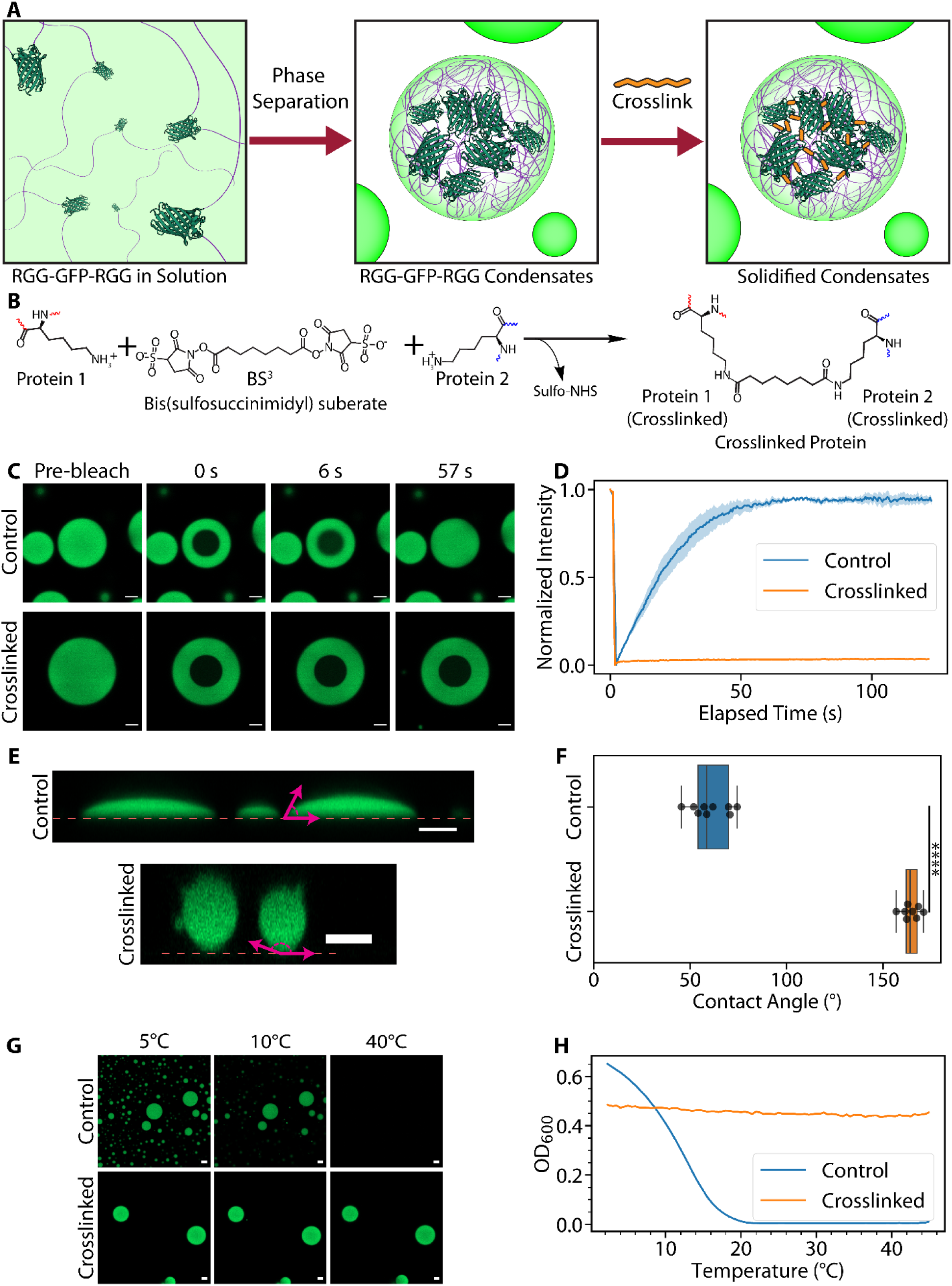
Crosslinking biomolecular condensates results in solidified protein-based microparticles. (A) General scheme for synthesizing protein-based microparticles. Through genetic engineering, two RGG domains (purple lines) are fused to GFP (dark green, PDB: 2Y0G) to form the fusion protein RGG-GFP-RGG. Phase separation in RGG-GFP-RGG solutions is induced by cooling to form biomolecular condensates. The proteins within the condensates can then be polymerized with a crosslinker (orange) to form solid particles. (B) Diagram of crosslinking reaction by BS^3^. (C,D) FRAP experiment. (C) Representative microscopy images from FRAP experiments on RGG-GFP-RGG condensates (control vs. crosslinked) before photobleaching and various timepoints post-bleach. (D) FRAP recovery curve of control (untreated) and crosslinked RGG-GFP-RGG condensates. Control condensates show fluorescence recovery after photobleaching, while crosslinked condensates show negligible recovery. Shaded region represents SEM; for crosslinked condensates, the SEM is too small to visualize on this plot. N = 3 condensates per condition. (E,F) Wetting experiment. (E) Side profiles of control and crosslinked condensates on glass surfaces. The dashed line represents the glass surface, and the magenta arrows show representative contact angle measurements. Side profiles were generated by confocal microscopy z-stacks. (F) Box plots showing contact angle measurements of control and crosslinked RGG-GFP-RGG condensates. **** = p < 0.0001 when compared by an independent two-way T-test, t = −27.889, N_Control_ = 9 contact angle measurements, N_Crosslinked_ = 8 contact angle measurements. (G,H) Assays of thermal responsiveness. (G) Representative microscopy images of control and crosslinked RGG-GFP-RGG at 5, 10, and 40 °C. (H) Representative plot of turbidity assay results. The control RGG-GFP-RGG solution exhibited temperature-dependent turbidity consistent with UCST phase behavior, and the crosslinked RGG-GFP-RGG sample displayed temperature-independent turbidity. Scale bar = 5 µm for all microscopy images.

## EXPERIMENTAL SECTION

### Cloning

Double-stranded DNA fragments for genes of interest were purchased from Genewiz. Genes of interest were cloned into pBamUK vectors (modified pET-29a(+) vectors with deletions of the *rop* and *bom* sequences) in frame with the C-terminal 6xHis-tag using the NEBuilder® HiFi DNA Assembly Master Mix (New England Biolabs). All plasmids contained kanamycin resistance genes. These plasmids were transformed into *Escherichia coli* NEB-5α (New England Biolabs) by heat shock following the manufacturer’s protocol and plated on lysogeny broth (LB) agar plates containing 50 µg/mL kanamycin for selection. Colonies were picked from the plates with a 10 µL micropipette tip and cultured in 2 mL LB containing 50 µg/mL kanamycin at 37 °C for 16 h (overnight) with shaking at 250 rpm. Plasmid DNA was isolated using the Monarch® Plasmid Miniprep Kit (New England Biolabs). The purified plasmids were then sent to Genewiz for Sanger sequencing to confirm the insertion of the correct gene sequences.

### Bacterial Transformation and Protein Expression

Chemically competent *E. coli* BL21(DE3) (New England Biolabs) were transformed by heat shock following the manufacturer’s protocol and plated on LB agar plates containing 50 µg/mL kanamycin for selection. Colonies were picked from the plates with a 10 µL micropipette tip and cultured in 5 mL LB containing 50 µg/mL kanamycin at 37 °C for 16 h (overnight) with shaking at 250 rpm. The starter culture was used to inoculate 500 mL of Terrific Broth (TB) in a 2 L baffled flask. For BsADH constructs, the TB was supplemented with 0.625 mM ZnSO4_4_ to facilitate cofactor incorporation into the enzyme structure. The cultures were shaken at 250 rpm and incubated at 37 °C until the optical density at 600 nm (OD_600_) reached 0.6-0.8, at which point protein expression was induced by the addition of 500 µM isopropyl-ß-D-1-thiogalactopyranoside (IPTG). Then the cultures were incubated at 18 °C and shaken overnight at 250 rpm.

### Protein Purification

Cells were collected by centrifuging cultures at 4,121×g for 25 min at 4 °C. The supernatant was decanted and then the cell pellet was resuspended in either a 50 mM sodium phosphate, 500 mM NaCl, 10% (v/v) glycerol, 10 mM 2-mercaptoethanol, 20 mM imidazole pH 8.0 buffer (for BsADH constructs) or a 1.5 M NaCl, 20 mM Tris, 20 mM imidazole, pH 7.5 buffer (all other proteins). EDTA-Free Pierce Protease Inhibitor (Thermo Fisher Scientific) was added to the resuspended cell solution and the cells were lysed by sonication in an ice water bath for 20 min (50% amplitude, 5 s on/off pulse). The crude extracts were centrifuged at 25,000 ×g for 30 min to separate soluble cell components from insoluble ones. The supernatants were filtered through 0.22 µm SteriFlip vacuum filters (Millipore Sigma) in preparation for fast protein liquid chromatography (FPLC).

### FPLC

FPLC was performed on the filtered supernatants using an Äkta Pure Purification system (Cytiva) with a 1 mL HisTrap HP column (Cytiva). For BsADH samples, mobile phase A consisted of a 50 mM sodium phosphate, 500 mM NaCl, 10% (v/v) glycerol, 10 mM 2-mercaptoethanol, 20 mM imidazole, pH 8.0 buffer and mobile phase B consisted of a 50 mM sodium phosphate, 500 mM NaCl, 10% (v/v) glycerol, 10 mM 2-mercaptoethanol, 500 mM imidazole, pH 8.0 buffer. For all other samples, mobile phase A consisted of 500 mM NaCl, 20 mM Tris, 20 mM imidazole, pH 7.5 buffer, and mobile phase B consisted of 500 mM NaCl, 20 mM Tris, 500 mM imidazole, pH 7.5 buffer. The column was equilibrated with 5 column volumes (CV) of mobile phase A. Sample was pumped into the column and the flowthrough was collected for downstream analysis. The column was washed with 5 CV of mobile phase A. Using a linear gradient elution, proteins of interest were eluted from the column where the mobile phase composition changed from 0% mobile phase B to 100% over 25 CV.

### Buffer Exchange

For RGG-GFP-RGG, and RGG-SpyCatcher-RGG, SYNZIP2-GFP, and SpyTagged GFP, the FPLC eluates were buffer exchanged overnight into a 150 mM NaCl, 50 mM borate, pH 8.0 buffer by dialysis using a Slide-A-Lyzer™ Dialysis Cassette G2, 10K MWCO (Thermo Fisher Scientific). The dialysis buffer was changed every two hours at room temperature. After the third dialysis buffer change (total 6 hours elapsed), the cassette was left to dialyze overnight at 45 °C to prevent proteins from adhering to the membrane. RGG-RGG was dialyzed into 150 mM NaCl, 20 mM Tris, pH 7.5 buffer for long-term storage, then buffer was exchanged to 150 mM NaCl, 50 mM borate, pH 8.0 with a desalting spin column (Zeba 7k MWCO column, Thermo Scientific). For MBP-GFP-RGG, protein was stored in buffer containing 500 mM NaCl. For BsADH-RGG eluates, we utilized a liquid-liquid fractionation technique to buffer exchange our samples. By storing BsADH-RGG eluates overnight at 4 °C, the BsADH-RGG phase separated to form an aqueous two-phase system with a protein-rich lower phase. The aqueous upper phase was then decanted by pipette and replaced with a 50 mM sodium phosphate, 10 mM 2-mercaptoethanol, pH 8.0 buffer. By doing so, we aimed to minimize protein loss from buffer exchange and enzyme denaturation. BsADH and BsADH-SpyTag were buffer exchanged using a PD-10 desalting column (Cytiva) into a 50 mM sodium phosphate, 10 mM 2-mercaptoethanol, pH 8.0 buffer. Glycerol was then added to the BsADH and BsaDH-SpyTag solutions to a final glycerol concentration of 50% v/v, and the samples were stored in −20°C.

### Protein Concentration Quantitation

Following buffer exchange, protein concentration was determined by BCA assay (Thermo Fisher) for BsADH fusion constructs, or UV absorption at 280 nm wavelength using a NanoDrop One spectrophotometer (Thermo Fisher Scientific) for all other proteins.

### General Crosslinking Procedure

RGG fusion protein stocks stored at 4 °C were first incubated in a 45 °C water bath to desorb them from the microcentrifuge tube walls. Protein stocks were diluted to a pre-determined concentration (2-100 µM) in their respective buffers, and then phase separation was induced in samples by placing tubes on ice. For BsADH-RGG, 50% w/v polyethylene glycol (PEG)-8K (Research Products International) dissolved in 50 mM sodium phosphate, 10 mM 2-mercaptoethanol, pH 8.0 buffer was added to the sample to a final concentration of 5% w/v PEG-8K. For core-shell condensates, solutions consisted of 5 µM RGG-RGG and 1 µM MBP-GFP-RGG. While the protein solution was incubated on ice for a pre-determined length of time (0-2 h), bissulfosuccinimidyl suberate (BS^3^; Thermo Fisher Scientific) was dissolved in a buffer similar to that of the protein solution to form a 100 mM BS^3^ stock solution. After the incubation time elapsed, the 100 mM BS^3^ stock solution was added to the sample to a final BS^3^ concentration of 20-fold molar excess of the protein concentration, according to the manufacturer’s protocol. The mixture was incubated on ice for 2 hours, then the crosslinking reaction was quenched by adding 1 M Tris HCl, pH 7.5 to a final concentration of 20 mM Tris. The quenched reaction was incubated at room temperature for 15 min. Crosslinked condensates were subsequently collected by centrifugation at 21,300 ×g for 5 min and the supernatant was decanted by pipetting. For BsADH-RGG samples, the supernatant was collected to measure enzyme immobilization yield by SDS-PAGE. Then the crosslinked condensates were washed three times by adding the appropriate buffer solution (150 mM NaCl, 50 mM borate, pH 8.0 buffer for RGG-GFP-RGG, RGG-SpyCatcher-RGG, and RGG-RGG/MBP-GFP-RGG, or 50 mM sodium phosphate, 10 mM 2-mercaptoethanol, pH 8.0 buffer for BsADH-RGG), centrifuging the sample at 21,300 ×g for 1 min, and decanting the supernatant by pipette. After washing, the crosslinked condensates were resuspended in their respective buffers to their original volume.

### Microscopy

Microscopy images of samples were taken with a Zeiss Axio Observer 7 inverted microscope with an Axiocam 702 monochrome sCMOS camera for widefield imaging and an LSM 900 confocal module for fluorescence imaging using a 63x/1.4 NA plan-apochromatic, oil-immersion objective (Carl Zeiss GmbH). GFP was excited to fluoresce with a 488 nm laser. Rhodamine and red FluoSpheres™ polystyrene microspheres (Thermo Fisher Scientific) were excited with a 561 nm laser. Transmitted light images were collected using a 0.55 NA condenser and an ESID module. Samples were plated in a 16-well glass-bottom dish with #1.5 glass thickness (Grace Bio-Labs) that were pretreated with a solution of 5% Pluronic ® F-127 (Millipore Sigma) for a minimum of 10 minutes, unless otherwise specified. The Pluronic ® F-127 was removed by pipette and the samples were added to the wells without washing with water, unless otherwise specified, to ensure that samples would not wet the glass surface.

For images taken at non-room temperature conditions (above or below 18-20 °C), a CherryTemp temperature controller (Cherry Biotech) was used to control the microscope stage temperature.

### Fluorescence Recovery After Photobleaching (FRAP)

FRAP experiments were performed on the previously mentioned Zeiss Axio Observer 7 inverted microscope with an LSM 900 confocal module. Bleaching was performed in a circular region of the condensate with a 488 nm laser for GFP or a 561 nm laser for rhodamine/TRITC-dextrans. Images were captured post-bleach with a 488 nm and/or 561 nm laser using a 63x/1.4 NA plan-apochromatic, oil-immersion objective.

### Contact Angle Measurements

Untreated and crosslinked RGG-GFP-RGG samples were plated on a 16-well glass-bottom dish with #1.5 glass thickness (Grace Bio-Labs) without any Pluronic ® F-127 treatment. Z-stack images were taken with the previously mentioned Zeiss Axio Observer 7 inverted microscope and LSM 900 confocal module. Using these Z-stack images, orthogonal projections were created within Zen Blue 3.0 (Carl Zeiss GmbH) to generate XZ and YZ side profiles. Contact angles of the condensates with the glass surface were drawn manually within the Zen Blue software. Control and crosslinked condensate contact angle measurements were compared with a two-way, independent T-test using *ttest_ind* from Python’s SciPy 1.15.2 library.

### Turbidity Assays

Turbidity assays were performed in a Cary 3500 Multicell UV-Vis Spectrophotometer (Agilent). Quartz cuvettes with 1 cm path length (Thorlabs) were first filled with buffer, allowed to equilibrate at 60 °C for 5 min, and the instrument was blanked. The buffer was then discarded, and the samples were added to the cuvettes. The cuvettes were allowed to equilibrate again at 60 °C for 5 min. Samples were cooled at a rate of 1 °C/min, with absorbance measured at 600 nm wavelength every 0.5 °C until the samples reached 4 °C.

### Particle Size Determination by Microscopy Image Analysis

A 384-well, glass bottom plate was pre-treated with 5% Pluronic ® F-127 for at least 10 min before the Pluronic ® F-127 was removed from the wells by pipette and each well was rinsed with distilled water. RGG-GFP-RGG protein stocks were diluted to a concentration of 2-100 µM with 150 mM NaCl, 50 mM borate, pH 8.5, and samples were placed on ice for 0, 1, or 2 h before 1 µL of BS^3^ stock solution was added to a final concentration in 20-fold molar excess of the protein concentration. Following the 2 h crosslinking period, the reaction was quenched with the addition of 1 M Tris HCl, pH 7.5 to a final concentration of 20 mM.

Z-stack images were collected by confocal microscopy. Maximum intensity projections were created from the z-stack images within Zen Blue 3.0. Particle sizes from the maximum intensity projections were determined in MATLAB 2024b using custom-written code that utilized a circular Hough transform to identify condensates and determine their size. Identified particles were visually confirmed and particles that were not identified by the MATLAB script were measured manually in ImageJ. From the number-weighted distribution obtained from image analysis, volume-weighted distributions for each group were derived by first calculating the volume of each particle and then dividing individual particle’s volume by the sum of all particle volumes to derive each particle’s weighting value. Letter-value plots^42^ of the volume-weighted particle size distributions were generated with the *catplot* function using the *boxen* option from the Seaborn 0.13.2 library in Python. To do so, the volume weight was first converted to an integer by multiplying it by a scale factor to create integer weights proportional to the volume distribution. Then a new DataFrame was created that repeated each diameter value based on the previously calculated integer weight, and this new DataFrame was used to create the letter-value plot.

### BS^3^ Concentration Range Experiments

RGG-GFP-RGG solutions with a protein concentration of 20 µM were incubated on ice in Pluronic ® F-127 coated tubes for 2 hours before a 100 mM BS^3^ stock solution was added to a final BS^3^ concentration of 5-fold to 200-fold molar excess (0.1 mM to 4 mM). The solutions were left unperturbed for two hours before the solution was quenched with 2 M Tris to a 20 mM final Tris concentration. Crosslinking yield was measured by performing SDS-PAGE on crosslinked sample alongside an uncrosslinked RGG-GFP-RGG solution that served as a control.

### Excitation/Emission Spectra Characterization

The excitation/emission spectra of RGG-GFP-RGG POMPOMS were measured with a SpectraMax M2 microplate reader (Molecular Devices, LLC) using the Fluorescence Intensity Read mode. For the excitation spectra, the emission wavelength was set to 575 nm and the plate reader swept a range of wavelengths from 350nm to 550 nm with an interval of 5 nm. For the emission spectra, the excitation wavelength was set to 375 nm and the plate reader swept a range of wavelengths from 455 to 700 nm with an interval of 5 nm.

### BS^3^ Crosslinking Kinetics

A 20 µM RGG-GFP-RGG solution was prepared in a tube coated with Pluronic ® F-127 for at least 30 minutes. Before the addition of the crosslinker, the protein solution was allowed to incubate on ice for 2 hours, and then a sample was drawn from the tube to serve as a pre-crosslinking control. A 100 mM BS^3^ stock solution was added to the RGG-GFP-RGG solution a final concentration of 20-fold molar excess (0.4 mM) and the tube was briefly vortexed before placing it back on ice. At various timepoints (1, 2, 5, 10, 15, 30, 45, 60, 75, 90, and 120 min after the crosslinker was added), the sample was briefly mixed by pipetting up and down before a sample of the solution was pipetted into another tube containing 2 M Tris, pH 7.5 to quench the reaction (final Tris concentration = 20 mM). Once samples from all timepoints had been collected, we performed SDS-PAGE, Coomassie staining, and densitometry to evaluate the progression of the crosslinking reaction by comparing the disappearance of the monomer band compared to the uncrosslinked control.

### Dextran and Polystyrene Particle Partitioning Experiments

Rhodamine B (Millipore Sigma) was added to condensate samples to a final concentration of 0.01 mg/mL. Tetramethylrhodamine isothiocyanate-dextrans (TRITC-dextrans) with average molecular weights (MWs) of 4,400 Da and 65,000-85,000 (Millipore Sigma) were added to condensate samples to a final concentration of 10 mg/mL. According to the manufacturer’s certificate of analysis, these TRITC-dextrans had a reported average molecular weight of 4,224 Da and 66,219 Da, respectively. Therefore, these TRITC-dextrans are referred to by the reported average molecular weights (TRITC-dextran 4.2K and TRITC-dextran 66.2K). After taking confocal fluorescence microscopy images, condensates were manually selected as regions of interest in Zen Blue 3.0 along with a region of the background. Because each condensate measured varied in sizes, we calculated the weighted average rhodamine fluorescence intensity within each condensates using the measured area and divided this value by the average background rhodamine fluorescence intensity to calculate the partition coefficient for each image. By weighing intra-condensate fluorescence by condensate size, we prevent smaller condensates from biasing the data. To test for statistical significance, the *anova_lm* and *pairwise_tukeyhsd* functions from Statsmodels 0.14.4 library in Python 3.13.0 was used to perform a two-way ANOVA along with a post-hoc Tukey’s range test. To estimate the hydrodynamic radius of dextrans and the pore size of our crosslinked condensates, we used data collected from Armstrong et al., 2004 on the measured hydrodynamic radii (*R_h_*) of various sized dextrans.^43^ In Microsoft Excel, the log(MW) was plotted on the x-axis and the measured *R_h_* was plotted on the y-axis. Then a 2^nd^ order polynomial equation was fit to the data, resulting in a line with an equation of y = 4.1337x^2^ – 7.1779x + 5.3298 and R^2^ = 0.9985. With this equation, the *R_h_* of a 66.2 kDa dextran was estimated to be 6 nm (or a hydrodynamic diameter of 12 nm).

Red fluorescent FluoSpheres™ polystyrene (PS) microsopheres (Thermo Fisher Scientific) with average diameters of 0.02 µm, 0.10 µm, and 0.50 µm were added to condensate samples to a final concentration of up to 1% volume fraction. Confocal fluorescence microscopy images were captured, and line profiles of each image were individually generated by a custom-written MATLAB code. For the representative microscopy images and the subsequent line profile analyses, a 1% volume fraction was used for 0.02 µm and 0.10 µm PS microspheres and a 0.1% volume fraction was used for the 0.50 µm microspheres. For the partition coefficient calculations, a volume fraction of 1% was used for all samples.

### Molecular Capture with Crosslinked RGG-SpyCatcher-RGG

Crosslinked RGG-Spycatcher-RGG condensates were collected by centrifugation at 21,300 ×g for 5 min. The buffer was decanted by pipette and the crosslinked condensates were resuspended in GFP solution (GFP-SYNZIP2, SpyTag-GFP, or GFP-SpyTag) of at least 50 µM concentration. The sample was incubated in a tube rotator spinning at 10 rpm for 30 min. After the incubation period, the crosslinked condensates were centrifuged again at 21,300 ×g for 5 min and the GFP solution was decanted by pipette. Subsequently, the condensates were washed in 150 mM NaCl, 50 mM borate, pH 8.0 buffer three times by resuspending the condensates, vortexing the sample to mix, centrifuging the sample at 21,300 ×g for 1 min, and decanting the supernatant by pipette.

Confocal microscopy images of the samples were collected. To calculate the enrichment ratio, we used an approach similar to the one used for calculating dextran partition coefficients. Crosslinked condensates were manually selected as regions of interest in Zen Blue 3.0 along with a region of the background. Because crosslinked condensates were of nonuniform size, we calculated the weighted average GFP fluorescence intensity within condensates using the measured area for weighting and divided this value by the average background fluorescence intensity to calculate the enrichment ratio. By doing so, we prevent smaller condensates from biasing the data.

To test for statistical significance, the *anova_lm* and *pairwise_tukeyhsd* functions from Python’s Statsmodels 0.14.4 library were used to conduct a two-way ANOVA along with a post-hoc Tukey’s range test.

### RGG-SpyCatcher-RGG POMPOMS Binding Capacity

The effect of BS^3^ on RGG-SpyCatcher-RGG POMPOMS binding capacity was tested by crosslinking 20 µM RGG-SpyCatcher-RGG solutions with BS^3^ concentrations ranging from 5-fold to 200-fold molar excess (0.1 to 4 mM). POMPOMS were collected and washed in a similar fashion as previously described. Before conjugation, the RGG-SpyCatcher-RGG POMPOMS were pelleted by centrifugation, and the buffer was removed by pipette. The RGG-SpyCatcher-RGG POMPOMS were resuspended in a solution containing a two-fold mole excess of GFP-SpyTag. The POMPOMS were collected by centrifugation and washed with buffer three times before resuspending in a 150 mM NaCl, 50 mM borate, pH 8.0 buffer. The enrichment ratio was calculated using a custom MATLAB script that measured the average fluorescence intensity inside identified condensates and divided it by the average fluorescence intensity outside condensates.

To test for statistical significance, the *anova_lm* and *pairwise_tukeyhsd* functions from Python’s Statsmodels 0.14.4 library were used to conduct a two-way ANOVA along with a post-hoc Tukey’s range test.

### SDS-PAGE

To prepare samples for sodium dodecyl sulfate-polyacrylamide gel electrophoresis (SDS-PAGE), 15 µL of samples were added to 5 µL of 4X NuPAGE™ LDS sample buffer (Thermo Fisher Scientific) and heated at 70 °C for 10 min using a T100 Thermal Cycler (Bio-Rad). After heating, 10 µL of each sample was loaded into the wells of a 1.0 mm, 15-well NuPAGE™ 4-12% Bis-Tris Mini protein gel (Thermo Fisher Scientific) along with a Novex™ Sharp pre-stained protein standard (Thermo Fisher Scientific). Gel electrophoresis was performed in a MES SDS running buffer at 200 V for 35 min, and the gel was stained with GelCode™ Blue Stain Reagent (Thermo Fisher Scientific) following the manufacturer’s protocol. Gel images were taken with an Azure 600 imager (Azure Biosystems) using an excitation wavelength of 685 nm and an emission wavelength of 735 nm.

### Crosslinking/Enzyme Immobilization Yield

AzureSpot (Azure Biosystems) was used to perform densitometry analysis on SDS-PAGE images for calculating RGG-GFP-RGG crosslinking yields and BsADH-RGG immobilization yields. SDS-PAGE was conducted on solutions before crosslinking alongside the supernatants of the crosslinked sample after centrifugation. Lanes were detected in AzureSpot automatically with some manual adjustments, and background was subtracted by selecting an image stripe over an empty lane. Bands corresponding to the molecular weight of the protein of interest were selected in AzureSpot. The crosslinking/enzyme immobilization yield was calculated with the followin equation:

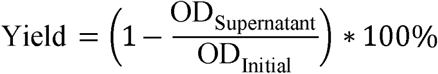

where OD_­_is the protein of interest’s band intensity in the supernatant (AU), and OD_­_is the protein of interest’s band intensity of the original solution (AU).

### BsADH-RGG Activity Assay

BsADH-RGG activity was assessed with a Cary 3500 Multicell UV-Vis Spectrophotometer (Agilent). Prior to the assay, enzymes were diluted in 50 mM sodium phosphate, pH 8.0, to create a working solution so that their ΔA_340_ measurements would be below 0.1 min^−1^ when 10 µL of the diluted enzyme were added to the reaction mix (a 1 mg/mL solution of free BsADH-RGG was typically diluted 200-fold to a 0.005 mg/mL working solution, and crosslinked BsADH-RGG was typically diluted 10- to 50-fold). The reaction mix contained 20 mM sodium phosphate, pH 8.0, 1 mM NAD^+^, 20 mM ethanol with a 600 µL final volume in a 1 cm quartz cuvette (Thorlabs). The reaction mix was heated to 60 °C in the spectrophotometer and the instrument was blanked. After blanking, 10 µL of enzyme or control sample (soluble BsADH-RGG, crosslinked BsADH-RGG, a blank 50 mM sodium phosphate buffer with no enzyme, or an RGG-GFP-RGG negative control) was added to the cuvettes for a final assay volume of 600 µL. The cuvettes were then capped and inverted several times before starting absorbance measurements at 340 nm (this wavelength corresponds to an absorption peak of NADH, which is produced as result of ADH activity). The BsADH-RGG and RGG-GFP-RGG samples were assayed alongside a blank cuvette containing the reaction mixture without any enzyme. The increase in absorbance was recorded for a total of 6 min, and linear regression was applied over the linear portion of the data (generally between minutes 1-6) using the Cary UV Workstation (Agilent) software to find the increase ΔA_340_, correlating to the increase in NADH concentration in solution. This increase in NADH was then normalized to the mass of protein in the cuvette. The specific activity was calculated with the following equation:

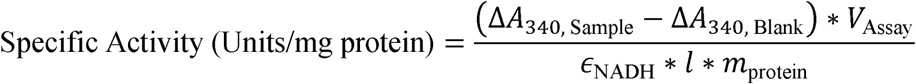

where Δ*A*_­_is the average change in 340 nm absorbance per minute,

*v*_­_is the total volume of the assay (mL),

*ɛ*_NADH_ is the millimolar extinction coefficient of NADH at 340 nm (6.22 mM^−1^ cm^−1^),

*l* is the path length of the cuvette (cm),

and *m*_­_is the mass of protein added to the assay (mg).

One unit of BsADH-RGG activity was defined as the conversion of 1 µmol NAD^+^ to NADH per minute under the reaction conditions. A derivation of this equation with further explanation for practical usage is included in the Supporting Information (Supplementary Notes 1, 2).

### Immobilization of BsADH-SpyTag by Capture Into RGG-SpyCatcher-RGG POMPOMS

BsADH-SpyTag was added to RGG-SpyCatcher-RGG POMPOMS at mole ratios of 0.2:1, 0.4:1, 0.6:1, 0.8:1, or 1:1 BsADH-Spytag to RGG-SpyCatcher-RGG. The solution was incubated at room temperature for 30 min to allow the enzyme to conjugate to the POMPOMS. The POMPOMS were collected by centrifugation at 21,300 x g for 5 min, the supernatant was decanted by pipette, and the POMPOMS were washed by resuspending in 50 mM sodium phosphate. This was repeated three times before resuspending them again in 50 mM sodium phosphate. The supernatant decanted from the first round of centrifugation was saved for SDS-PAGE and used to analyze the immobilization yield by comparing the band intensity to a sample with a similar initial concentration of enzyme. The specific activity was then measured and calculated in a similar manner to the previously mentioned BsADH-RGG assays.

### Thermal Stability Assays

BsADH and BsADH-SpyTag glycerol stocks were diluted to a 1 mg/mL protein concentration in 50 mM sodium phosphate, which were then pipetted into quartz cuvettes with a path length of 1 cm (Thorlabs) along with a cuvette filled with only 50 mM sodium phosphate that was used as a blank reference. The cuvettes were placed in a Cary 3500 Multicell UV-Vis Spectrophotometer (Agilent) and allowed to equilibrate to 50 °C. Samples were heated at a rate of 0.5°C/min until the samples reached 90 °C. As the protein sample is heated, changes in secondary and tertiary structure exposes the hydrophobic core and these residues absorb more UV light. Therefore, we measured the absorbance at 280 nm at every 0.1°C interval to determine the thermal stability of our protein samples. A plot of the first order derivative was used to determine the melting temperature of the enzyme. The apparent melting temperature (T_m_) is defined as the temperature at which half of the protein is still in its soluble form.

### Data Visualizations

Data visualizations were created with the Seaborn 0.13.2 library in Python using the *lineplot*, *boxplot*, *barplot*, *swarmplot, scatterplot,* and *catplot* functions.

## RESULTS AND DISCUSSION

### Crosslinking Converts Dynamic Condensates into Solid Microparticles

We reasoned that LLPS of proteins would be useful for fabricating microscale materials because LLPS spontaneously generates spherical, protein-rich droplets. Protein droplets have promising potential industrial applications, but the liquid-like and ephemeral nature of droplets make them ill-suited for many downstream applications. Consequently, we sought to convert protein droplets from liquid to solid by crosslinking the protein monomers that form these condensates. To achieve this, we used the homobifunctional crosslinker bissulfosuccinimidyl suberate (BS^3^), which reacts with primary amines such as lysine sidechains and the N-termini of proteins under neutral to basic conditions, releasing sulfo-NHS in the process (Fig. 1B). This crosslinker was chosen for its commercial availability, moderately long spacer arm (11.4 A), and safety due to its membrane impermeability. We also required a crosslinker that would be highly water soluble to avoid using organic solvents that may perturb the phase behavior of the proteins. Therefore, BS^3^ was a suitable crosslinker for our application. For most of our experiments, except those testing BS^3^ concentration ranges, we added BS^3^ in 20-fold molar excess of protein concentration, which is commonly used in crosslinking experiments to maximize the extent of crosslinking reactions.

After crosslinking RGG-GFP-RGG droplets with BS^3^, the condensates exhibited a dramatic shift in material properties. Uncrosslinked condensates displayed rapid molecular diffusion of the constituent proteins: Fluorescence recovery after photobleaching (FRAP) showed >90% recovery within 1 minute (Fig. 1C, D). In contrast, crosslinked condensates showed negligible recovery (<5%) in FRAP experiments. This minimal fluorescence recovery likely arose from a small amount of uncrosslinked RGG-GFP-RGG remaining in the dilute phase that diffused into the bleached regions (Fig. 1D). Crosslinked and uncrosslinked condensates were further differentiated by their wetting behavior on untreated glass surfaces (Fig. 1E). RGG-GFP-RGG condensates displayed low contact angles on untreated glass surfaces (average = 60.4°; Fig. 1E, F), indicating surface wetting due to the condensates’ liquid-like nature and adhesion to the glass. On the contrary, crosslinked condensates had significantly higher contact angles (average = 164.4°, p < 0.0001), indicating a loss of wetting capacity because of condensate solidification (Fig. 1E, F). Additionally, uncrosslinked RGG-GFP-RGG exhibits upper critical solution temperature (UCST) phase behavior: at lower temperatures, the proteins condense to form liquid droplets, and at higher temperatures, the proteins dissolve back into solution to form a single phase. This UCST phase behavior is observable either by microscopy (Fig. 1G) or by measuring the turbidity at different temperatures with spectrophotometry (Fig. 1H). A 20 µM solution of RGG-GFP-RGG (in 150 mM NaCl, 50 mM borate, pH 8.0 buffer) forms condensates below 20 °C but dissolves at higher temperatures (Fig. 1G, H). On the other hand, crosslinked condensates remained intact with spherical morphology across a broad temperature range (Fig. 1G, H). Collectively, these data indicate that BS^3^ crosslinking converted the liquid-like RGG-GFP-RGG condensates into solid-like microparticles with enhanced thermal stability. Henceforth, we will refer to these crosslinked biomolecular condensates as POMPOMS.

### Tuning POMPOMS Sizes via Pre-Crosslinking Parameters

Particles of differing sizes may be desirable to users depending on the application. In the case of molecular capture, smaller particles would provide an increased surface binding capacity due to a higher surface-area-to-volume ratio. For enzyme immobilization, smaller particles would allow for higher enzyme loading (for surface immobilized enzymes) and shorter diffusion distances for reactants (for internally immobilized enzymes). On the other hand, for batch chemical reactions in manufacturing contexts, larger immobilized enzyme particles would be easier to remove from the reaction mixture and recover for reuse through methods such as centrifugation or filtration. In flow chemistry, the mechanical stability of smaller particles would allow them to retain integrity under high pressure flows, while larger particles would allow for higher volumetric flow rates when working with viscous solutions. Therefore, we sought to control the size of our POMPOMS by tuning pre-crosslinking conditions.

In nature, cells can regulate condensate size through various mechanisms, including protein expression level (concentration) relative to the saturation concentration. Droplets can also grow over time due to coalescence as well as Ostwald ripening. We thus rationalized that protein concentration and coalescence time before crosslinking were two simple parameters that could be used to control the resultant POMPOMS sizes. We varied protein concentration from 2 to 100 µM, and we varied the time before crosslinking from 0 to 2 h. Based on confocal microscopy and image analysis, we observed that particle size increased with both incubation time and protein concentration, shifting the POMPOMS size distribution towards larger diameters (Fig. 2A, 2B). The matrix of conditions resulted in a range of particle sizes, from smaller than 1 µm to larger than 40 µm (Fig. 2A,B). Increases in one or both parameters resulted in increases in the de Brouckere mean diameters (the volume-weighted mean diameter; Fig. 2B inset). For subsequent experiments, we standardized pre-crosslinking conditions to maximize POMPOMS size with the least concentrated protein solution (20 µM RGG-GFP-RGG, 2 hours coalescence time).

**Figure 2.**
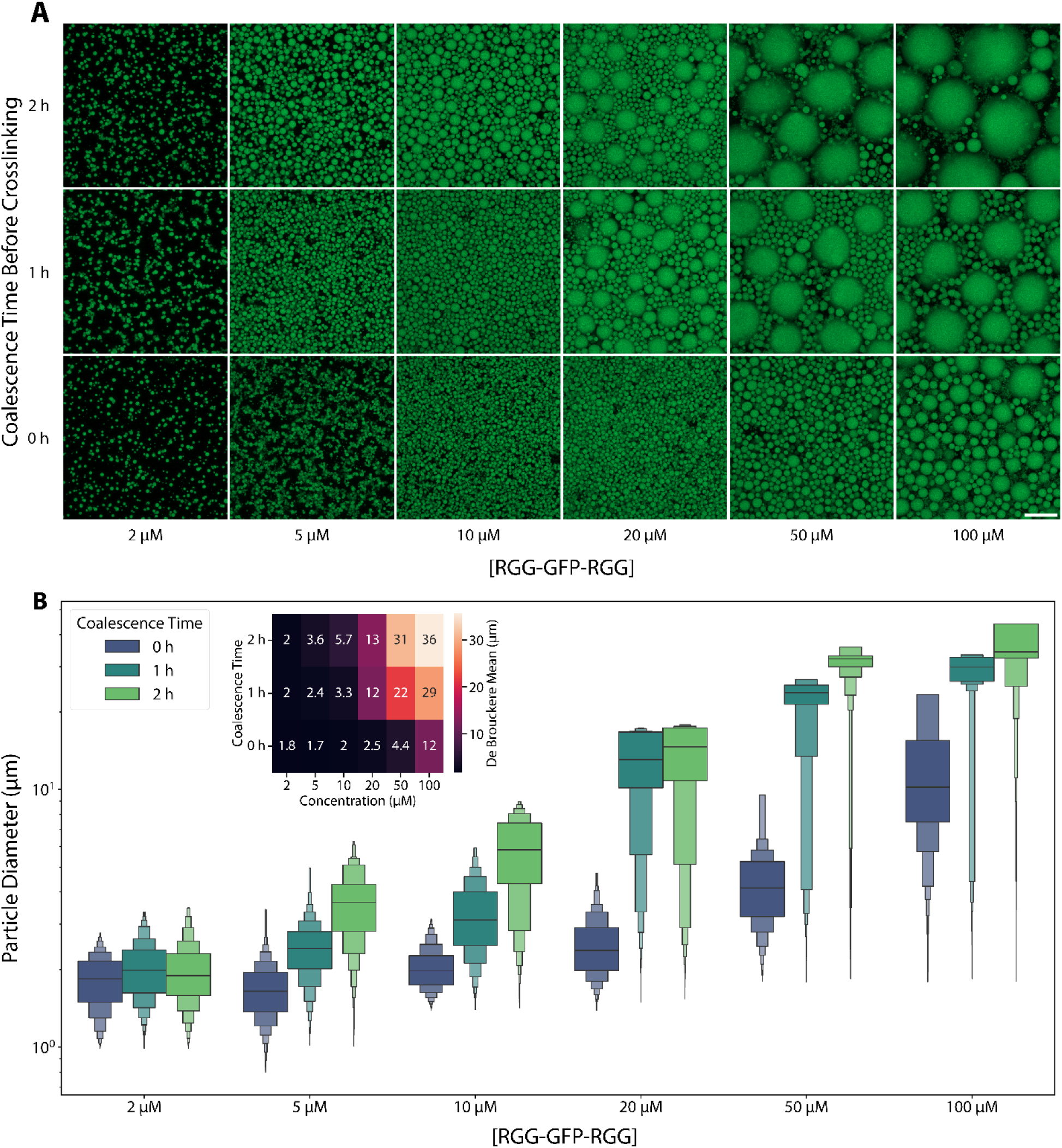
POMPOMS size can be modulated by varying protein concentration and allowing time for the condensates to coalesce before crosslinking. (A) Representative maximum intensity projection microscopy images of microparticles synthesized from solutions of 2-100 µM RGG-GFP-RGG that were allowed to coalesce for 0-2 h before crosslinker was added. Scale bar = 20 µm. (B) Letter-value plots for volume-weighted particle size distributions measured by image analysis. The center lines represent the median and the first two sections surrounding the median each represent 25% of the data. Each successive level outwards contains half of the data of the preceding section. Inset: heat map showing the de Brouckere mean (volume-weighted mean) diameters for each group.

### Effect of BS^3^ Crosslinker Concentration

So far, we have demonstrated crosslinking of condensates using a fixed 20-fold molar excess of BS^3^ relative to RGG-GFP-RGG. We next asked how crosslinker concentration affects POMPOM formation and properties. To answer this, we prepared batches of POMPOMS using BS^3^ at 5-fold to 200-fold molar excess relative to RGG-GFP-RGG (0.1 mM to 4 mM BS^3^ added to 20 µM RGG-GFP-RGG). First, we assessed the overall crosslinking yield based on the disappearance of the monomer band in SDS-PAGE. Crosslinking yield increased with BS^3^ concentration, but BS^3^ concentrations exceeding 10-fold molar excess resulted in marginal increases in yield (Fig. 3A, Supp. Fig. 1). BS^3^ concentration did not substantially affect the excitation/emission spectra of RGG-GFP-RGG POMPOMS compared to uncrosslinked RGG-GFP-RGG (Fig. 3B), suggesting that the crosslinker is not heavily distorting the GFP structure. Next, we analyzed how crosslinker concentration affects POMPOMS physical properties. Uncrosslinked RGG-GFP-RGG condensates wet untreated glass surfaces, whereas crosslinked POMPOMS do not. Even using 5-fold molar excess of BS^3^, the POMPOMS seem to be fully crosslinked, showing contact angles well above 90° on untreated glass surfaces (Fig. 3C). Additionally, in FRAP experiments, even with 5-fold molar excess crosslinker, the photobleached regions maintain sharp boundaries over time (Supp. Fig. 2). Together, these results suggest that even 5-fold molar excess BS^3^ is sufficient to generate stable POMPOMS containing a percolated network of chemically crosslinked proteins, albeit with a reduced crosslinking yield compared to higher crosslinker concentrations.

**Figure 3.**
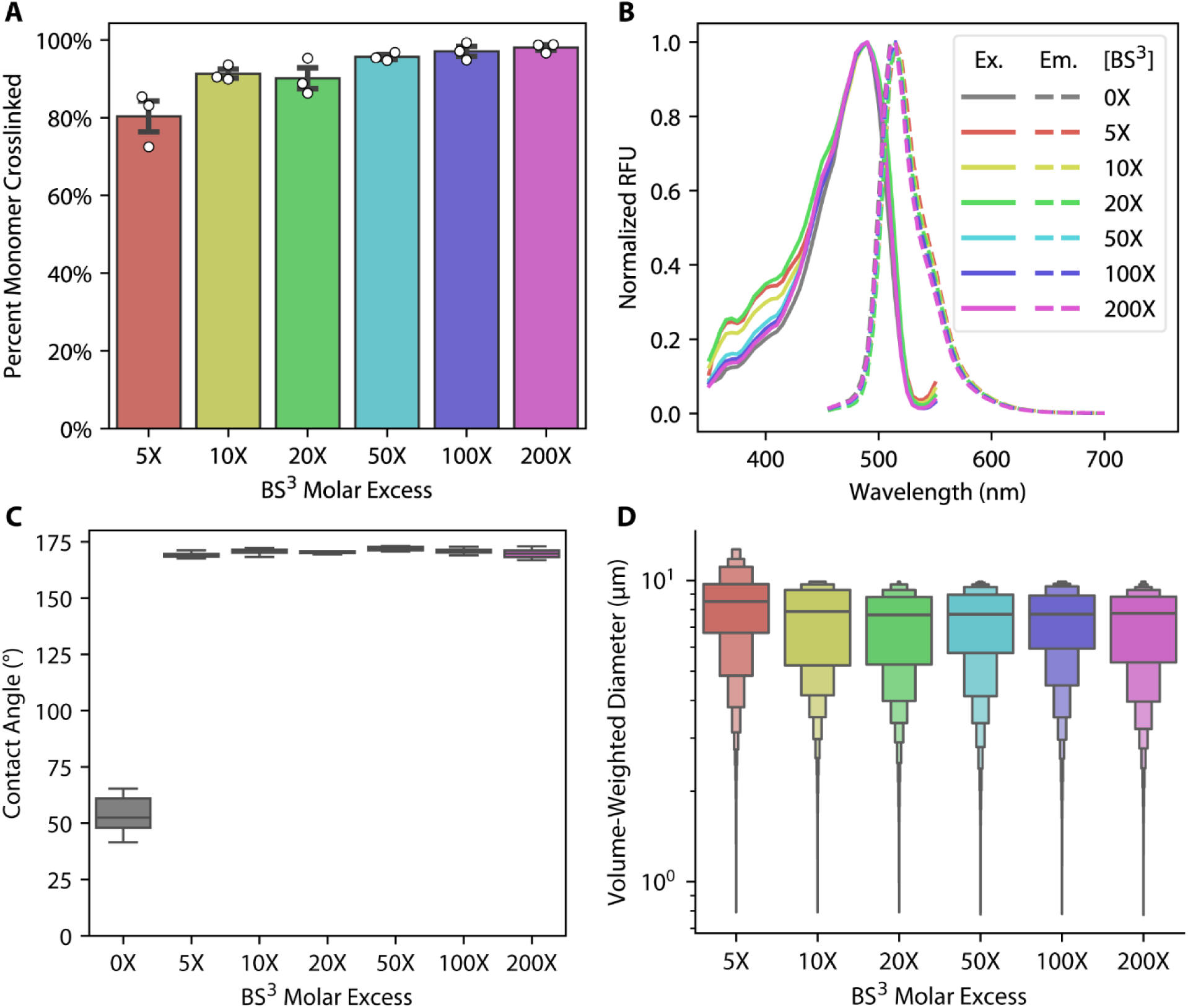
Effects of BS^3^ concentration on overall crosslinking yield, optical properties, contact angle, and POMPOMS size. (A) Overall crosslinking yield of RGG-GFP-RGG batches crosslinked with different BS^3^ concentrations as measured by SDS-PAGE densitometry analysis. Error bars represent SEM, N = 3 batches for each concentration. (B) Fluorescence excitation/emission spectra of uncrosslinked RGG-GFP-RGG ([BS^3^] = 0X) compared with spectra of RGG-GFP-RGG POMPOMS crosslinked with 5-200X molar excess BS^3^. Solid lines denote excitation spectra and dashed lines represent emission spectra. (C) Contact angle measurements comparing uncrosslinked RGG-GFP-RGG (0X) with RGG-GFP-RGG POMPOMS crosslinked with 5-200X molar excess BS^3^. N = 3 batches for each condition. (D) Letter-value plots for volume-weighted particle size distributions of RGG-GFP-RGG POMPOMS crosslinked with different concentrations of BS^3^. The particle diameters were measured by confocal microscopy Z-stacks coupled with image analysis. The center lines represent the median and the first two sections surrounding the median each represent 25% of the data. Each successive level outwards contains half of the data of the preceding section.

Interestingly, the particle size distributions of RGG-GFP-RGG POMPOMS did not differ greatly with BS^3^ concentration (Fig. 3D). The weak influence of crosslinker concentration on POMPOMS diameter may be the result of BS^3^’s fast reaction kinetics when crosslinking condensed RGG-GFP-RGG. SDS-PAGE densitometry of RGG-GFP-RGG solutions crosslinked using a 20-fold molar excess BS^3^ solution reveals that, on average, 80% of the monomer is crosslinked within 5 minutes and the crosslinking reaction reaches its maximum extent between 30 to 45 minutes (Supp. Fig. 3). The reaction occurs quickly enough that we observed instances of condensates that solidified mid-fusion, even though droplet fusion for RGG-GFP-RGG often occurs in the span of seconds (Supp. Fig. 4). Overall, these experiments suggest that the BS^3^ crosslinking reaction is highly efficient at crosslinking biomolecular condensates and is not disruptive to the fused folded protein domain’s structure.

### Condensate Network Architecture is Preserved Post-Crosslinking

Previous research has established that proteins within biomolecular condensates create a dynamic network that permits diffusion and even enrichment of cargo molecules, depending on a complex interplay of factors including cargo size and molecular interactions with the condensate’s scaffold.^38,44^ Because the overall morphology of condensates was maintained after crosslinking (Fig. 1C, Fig 2A, Supp. Fig. 2, Supp. Fig. 4), we surmised that this network structure might also be preserved in our POMPOMS upon crosslinking. To assess the POMPOMS’ molecular permeability, we observed the partitioning of a fluorescent small molecule (rhodamine) and fluorescently labeled macromolecules (TRITC-dextran 4.2K, TRITC-dextran 66.2K) using confocal microscopy. The largest of these probes, TRITC-dextran 66.2K, has an estimated hydrodynamic diameter of 12 nm.^43^ We found that these probes could partition into the interiors of RGG-GFP-RGG POMPOMS (Fig. 4A, B). The average partition coefficients were 30.6 for rhodamine B, 15.4 for TRITC-dextran 4.2K, and 9.1 for TRITC-dextran 66.2K, where a partition coefficient greater than 1 indicates enrichment inside POMPOMS as compared to the continuous phase (Fig. 4C). In accordance with previous research on uncrosslinked condensates, the partitioning inversely correlated with the probes’ molecular weight (Fig. 4C).^38^ The partition coefficients of the probes into crosslinked condensates did not significantly differ as compared to untreated condensates (p = 0.72, Supp. Fig. 5). These results confirm that the crosslinked condensates retain a porous internal structure.

**Figure 4.**
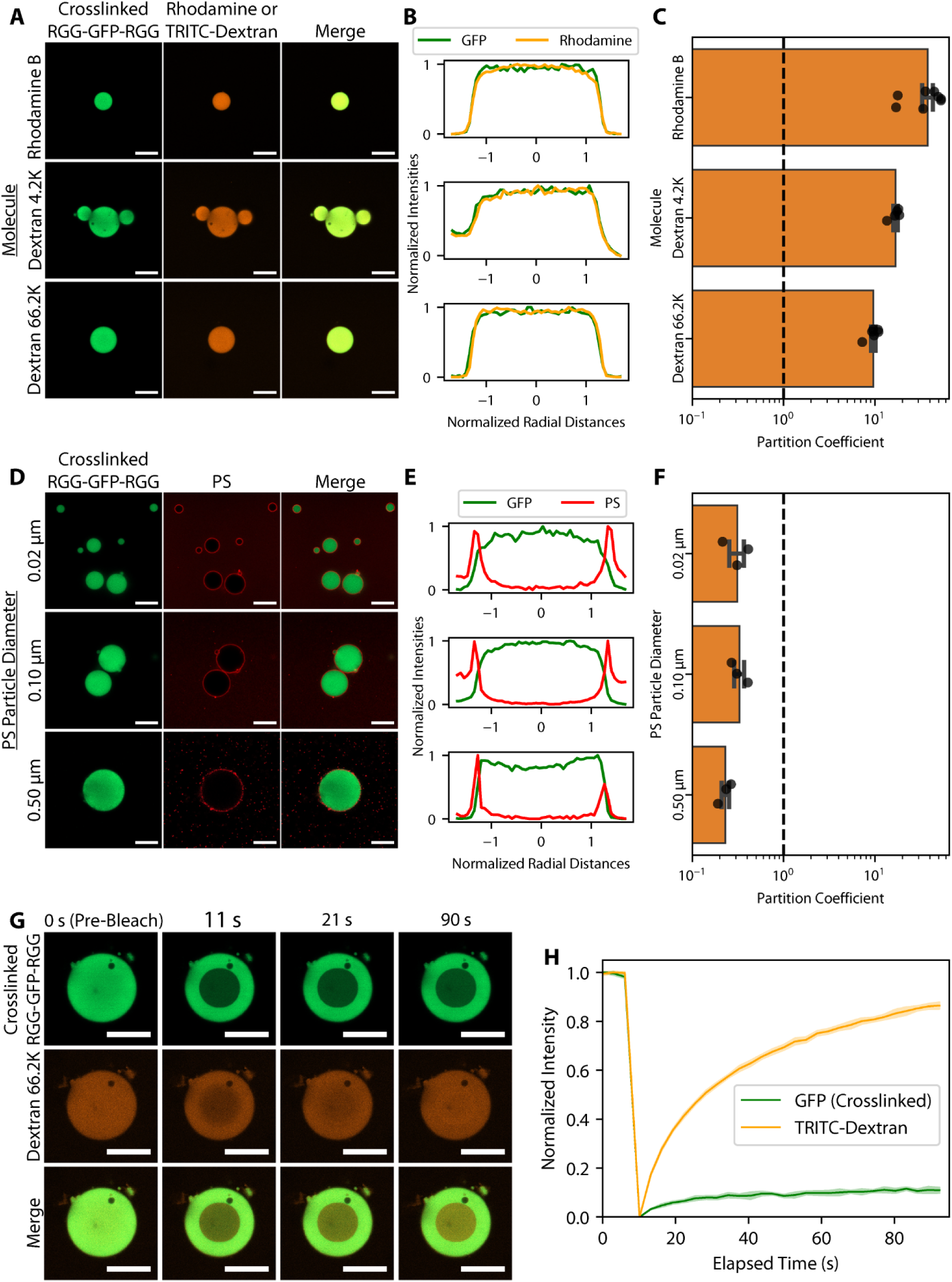
Crosslinked POMPOMS remain permeable to macromolecular clients and permit client diffusion. (A,B,C) Molecular partitioning experiments. (A) Representative microscopy images demonstrating colocalization of rhodamine B and TRITC-labeled dextrans inside RGG-GFP-RGG POMPOMS. (B) Normalized line profiles of condensates corresponding to representative microscopy images displayed in panel A, showing overlap between GFP and rhodamine fluorescence signals. (C) Bar plot of partition coefficients observed for rhodamine and TRITC-labeled dextrans. The dashed line represents a partition coefficient value of 1. Partition coefficients > 1 indicate that these molecules are enriched inside POMPOMS. Error bars represent SEM. N_Rhodamine B_ = 8 images, N_Dextran 4.2K_ = 5 images, N_Dextran 66.2 K_ = 6 images. (D,E,F) Polystyrene particle partitioning experiments. (D) Representative microscopy images demonstrating exclusion of PS particles from RGG-GFP-RGG POMPOMS interiors. (E) Normalized line profiles of condensates corresponding to representative microscopy images displayed in panel D, showing little overlap between GFP and PS fluorescence signals. (F) Bar plot of partition coefficients of PS particles in RGG-GFP-RGG POMPOMS. The dashed line represents a partition coefficient value of 1. Partition coefficients < 1 indicate exclusion of PS particles. Error bars represent SEM. N = 3 for each particle size. (G,H) Intra-condensate FRAP experiment. (G) Representative microscopy images of RGG-GFP-RGG POMPOMS and TRITC-dextran 66.2K before photobleaching and various timepoints post-bleach. (H) FRAP recovery curve. TRITC-dextran 66.2K fluorescence recovered, whereas crosslinked RGG-GFP-RGG displayed only limited recovery. The shaded region represents SEM. For all microscopy images, scale bar = 20 µm.

To assess the upper size limit of probes that can enter POMPOMS, we mixed condensates with carboxylated polystyrene (PS) particles of varying sizes (0.02, 0.10, and 0.50 µm diameters) and observed the particles’ localization using confocal microscopy. Despite the large size of the PS particles, uncrosslinked RGG-GFP-RGG condensates can internalize PS particles due to the proteins’ ability to dynamically rearrange and because of PS particle surface interactions with the RGG-GFP-RGG proteins (Supp. Fig. 6).^44^ For POMPOMS, on the other hand, all measured sizes of PS particles were excluded from the condensates and instead localized to the condensate interface (Fig. 4D, E). Partition coefficients were less than 1, because fluorescence signal was lower in the condensate interior compared to the bulk solutions (Fig. 4F). The observations from both the molecular and PS particle partitioning suggest that RGG-GFP-RGG POMPOMS retain a porous network with an estimated pore size somewhere between 12 nm and 20 nm in diameter.

To investigate the dynamics of clients partitioned into POMPOMS, FRAP experiments were performed with TRITC-dextran 66.2K that had partitioned into crosslinked RGG-GFP-RGG (Fig. 4G). After photobleaching, the TRITC-dextran 66.2K recovered more than 80% of its fluorescence within 90 seconds (Fig. 4H). The POMPOMS were also photobleached, but showed negligible recovery, confirming that the TRITC-dextran 66.2K was colocalizing in a crosslinked condensate interior (Fig. 4G, H). The rapid recovery of TRITC-dextran 66.2K fluorescence in the interiors of POMPOMS indicate that the POMPOMS form porous scaffolds that allow for diffusion and dynamic exchange of sufficiently small client molecules within the POMPOMS.

### Targeted Molecular Capture into POMPOMS via SpyCatcher/SpyTag

One advantage of protein-based materials is that biomolecular interactions can be encoded in the material through genetic engineering, avoiding the need for additional functionalization steps. To demonstrate that we can encode functionality into POMPOMS, we engineered RGG-SpyCatcher-RGG POMPOMS to capture SpyTagged GFP from solution. The SpyCatcher protein can spontaneously conjugate with a SpyTagged protein by forming an isopeptide bond between a reactive lysine in the SpyCatcher domain and an aspartic acid in the SpyTag.^45,46^ RGG-SpyCatcher-RGG forms condensates that can be crosslinked into POMPOMS using BS^3^. After crosslinking, we tested the POMPOMS’ capture capabilities by incubating them with a protein solution containing GFP tagged at its N- or C-terminus with SpyTag (SpyTag-GFP or GFP-SpyTag, respectively) or with SYNZIP2 as a nonspecific interaction control (SYNZIPs are synthetic coiled coil that are not expected to interact specifically with SpyCatcher).^38,47^ Following incubation, we decanted the solutions of tagged GFP, washed the RGG-SpyCatcher-RGG POMPOMS, and then measured the GFP fluorescence inside the POMPOMS to compare how well the particles could capture SpyTagged GFP compared to the nonspecific control, SYNZIP2-GFP (Fig. 5A). SpyTagged GFP exhibited a >60-fold higher enrichment over the control (Fig 5B, C): mean SYNZIP2-GFP enrichment was only 8.9, whereas mean GFP-SpyTag enrichment was 598.1, and mean SpyTag-GFP enrichment was 1203.9. This difference was highly significant (one-way ANOVA: F = 98.53, p < 0.0001), and a post hoc Dunnett’s test confirmed significant differences between our SYNZIP2-GFP control and SpyTagged GFP (SYNZIP2-GFP vs. GFP-SpyTag: t = 6.925, p < 0.0001; SYNZIP2-GFP vs. SpyTag-GFP: t = 14.04, p < 0.0001). To form the isopeptide bond, the SpyCatcher domain relies on a lysine residue, which is susceptible to inactivation by reacting with amine-reactive cross-linkers such as BS^3^. We found that crosslinking RGG-SpyCatcher-RGG condensates with BS^3^ concentrations higher than 20X molar excess resulted in a substantial decrease in GFP-SpyTag binding capacity (Supp. Fig. 7). Together with our prior data (Fig. 3), this experiment suggests that a 20-fold BS^3^ molar excess permits good crosslinking yield without unduly compromising protein function. Overall, these results validated SpyCatcher-mediated capture of cargo into POMPOMS, demonstrating that specific biomolecular interactions can be encoded into these crosslinked protein-based materials.

**Figure 5.**
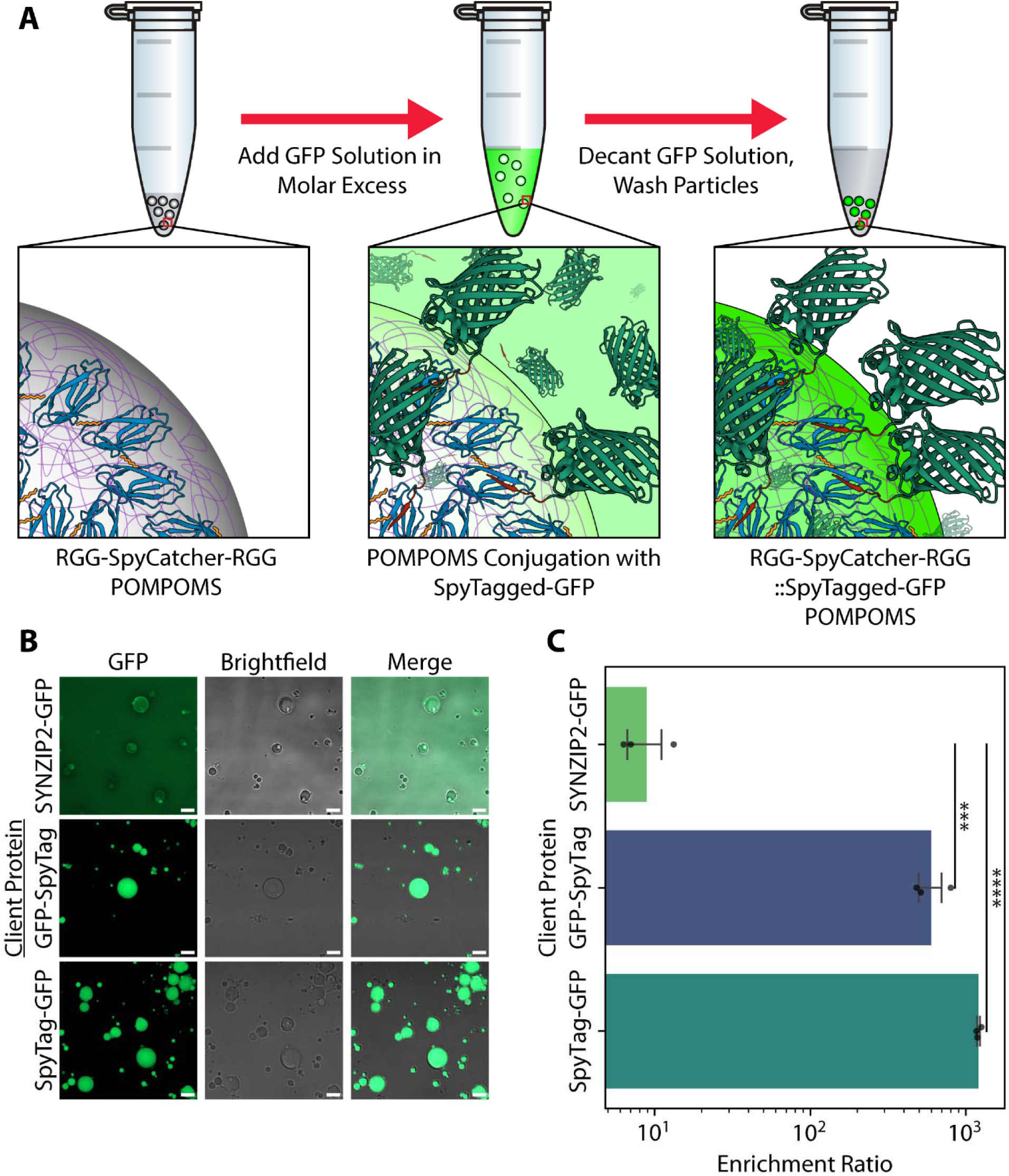
Specific interaction motifs can be encoded into POMPOMS to allow for biomolecular capture. (A) Scheme for RGG-SpyCatcher-RGG POMPOMS. POMPOMS were incubated for 30 min, collected by centrifugation, and washed several times to remove any weakly-binding GFP. GFP structure: PDB 2Y0G. SpyCatcher/SpyTag structure: PDB 4MLI. (B) Representative microscopy images of RGG-SpyCatcher-RGG POMPOMS after incubation with SYNZIP2-GFP, SpyTag-GFP, or GFP-SpyTag and after washing. Scale bar = 10 µm. (C) Bar plot comparing enrichment ratios of SYNZIP2-GFP to SpyTagged GFP. The difference between all groups was found to be statistically significant when compared using a one-way ANOVA (F = 98.6, p = 2.57*10^−5^, N = 9 images, 3 images per condition). When the non-specific binding control (SYNZIP2-GFP) was compared to each SpyTagged GFP by a post-hoc Dunnett’s test, the SpyTagged GFPs were found to have a significantly higher enrichment (GFP-SpyTag vs. SYNZIP2-GFP: t = 6.925, p = 6.72*10^−4^; SpyTag-GFP vs. SYNZIP2-GFP: t = 14.044, p = 1.50*10^−5^). Error bars represent SEM. *** = p < 0.001, **** = p < 0.0001.

### Hierarchical Material Design by Crosslinking Core-Shell Condensates

We previously demonstrated that a two-component system comprising a phase-separating protein and an amphiphilic protein can form core-shell condensates.^48^ In this system, the amphiphilic proteins coat the condensate surface and stabilize the liquid-liquid interface.^48^ We next aimed to integrate these amphiphilic proteins into our POMPOMS platform to generate microparticles with hierarchical structure. Specifically, we crosslinked condensates formed from the phase-separating core protein RGG-RGG with the amphiphilic protein MBP-GFP-RGG. MBP-GFP-RGG contains insoluble (RGG) and soluble (MBP; maltose binding protein) domains, so it forms a shell around a RGG-RGG condensate by adsorbing to the condensate surface.^48^ Confocal microscopy revealed that the crosslinked MBP-GFP-RGG + RGG-RGG POMPOMS maintained distinct core and shell phases, much like their uncrosslinked liquid counterparts (Fig. 6A). (Interestingly, crosslinked core-shell condensates exhibited increased fluorescence intensity at their surfaces. We speculate that adding crosslinker shifts the equilibrium to prevent MBP-GFP-RGG desorption.) We next confirmed that crosslinking the core-shell condensates indeed altered the thermal responsiveness and molecular diffusivity in the condensates. After incubating uncrosslinked condensates and core-shell POMPOMs at 50 °C for 20 min, we noticed that while the uncrosslinked condensates dissolved, the core-shell POMPOMS resisted dissolution (Fig. 6B). Similarly, FRAP experiments in which the MBP-GFP-RGG shell was photobleached showed that while the uncrosslinked protein layers exhibited moderate fluorescence recovery, the crosslinked POMPOMS layers exhibited no recovery (Fig. 6C, D). These data provide proof of concept that, by leveraging the intrinsic ability of amphiphilic proteins to adsorb at liquid-liquid interfaces, we can decorate the surface of POMPOMS – a feature that could be useful for a variety of purposes. For example, POMPOMS can be coated with a shell of stabilizers or antifouling proteins to protect the inner contents of the core.^49^ In the case of enzyme immobilization and biocatalysis, shell proteins could be used to anchor the immobilized enzyme inside reactors, orient surface-immobilized enzymes so that their active sites are more accessible to the bulk solvent, or spatially arrange multiple enzymes for cascade reactions. POMPOMS therefore hold promise as hierarchical, multifunctional, protein-based materials.

**Figure 6.**
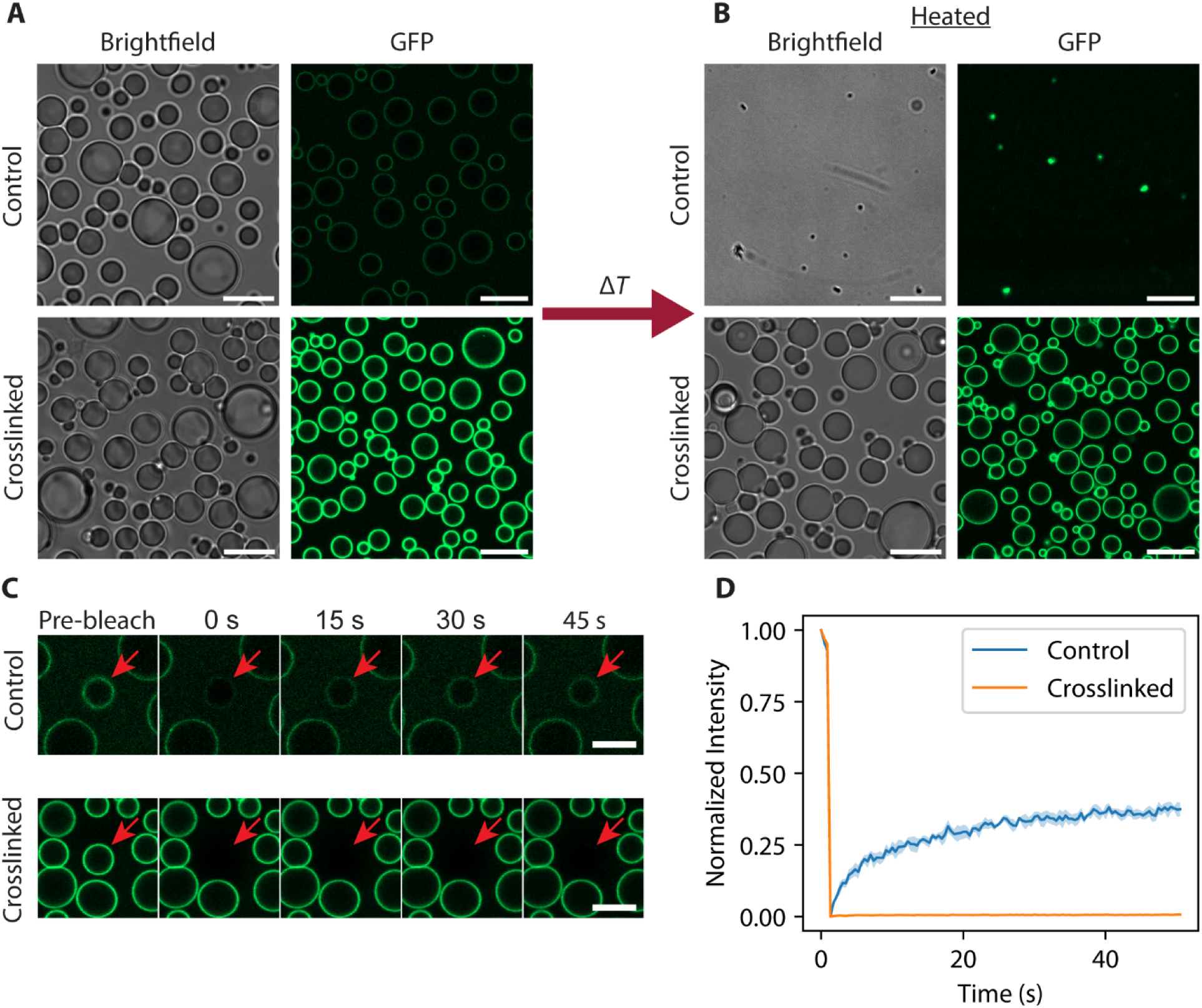
Crosslinking biphasic core-shell condensates results in POMPOMS with microarchitecture. (A) Representative microscopy images of untreated (control) and crosslinked solutions of 5 µM RGG-RGG and 1 µM MBP-GFP-RGG. Scale bars = 10 µm. (B) Representative microscopy images of 5 µM RGG-RGG and 1 µM MBP-GFP-RGG solutions (control and crosslinked) after heating for 20 min at 50 °C. Scale bar = 10 µm. (C) Representative microscopy images of core-shell condensates before and after photobleaching. Red arrow indicates the bleached condensate. Scale bar = 5 µm. (D) FRAP recovery curves. Control core-shell condensates show some recovery after photobleaching while crosslinked core-shell condensates show none. Shaded region represents SEM. N = 3 FRAP experiments for each condition.

### POMPOMS for Enzyme Immobilization

To demonstrate the potential usage of POMPOMS for enzyme immobilization, we crosslinked condensates incorporating an alcohol dehydrogenase from *Geobacillus stearothermophilus* (BsADH) fused to an RGG domain. BsADH is a homotetrameric, thermostable, zinc-dependent enzyme (Fig. 7A) that catalyzes the conversion of alcohols (such as ethanol) to their respective aldehyde, reducing an NAD^+^ cofactor to NADH in the process (Fig. 7B). This specific enzyme can be used as part of biocatalytic cascades for cofactor regeneration, and other alcohol dehydrogenases hold value for industrial biocatalytic processes.^50,51^ We appended a single RGG domain to each BsADH subunit (BsADH-RGG) and confirmed by microscopy that this construct could form condensates (Fig. 7C). We immobilized BsADH-RGG by crosslinking these biomolecular condensates using BS^3^. With our method, we achieved average immobilization yields of approximately 84%. We measured the specific activity of free (uncrosslinked) BsADH-RGG, crosslinked BsADH-RGG, and crosslinked RGG-GFP-RGG by measuring the concentrations of NADH through its absorbance of 340 nm wavelength light (Fig. 7D). The crosslinked RGG-GFP-RGG functioned as another negative control in addition to a blank solution without enzyme that was measured for each activity assay. (The activity of BsADH-RGG in its uncrosslinked condensate form was not measured since the high concentration of fusion enzyme needed for phase separation would require proportionally high concentrations of substrates/cofactors and would make it difficult to consistently obtain linear data for enzyme activity calculations.)

**Figure 7.**
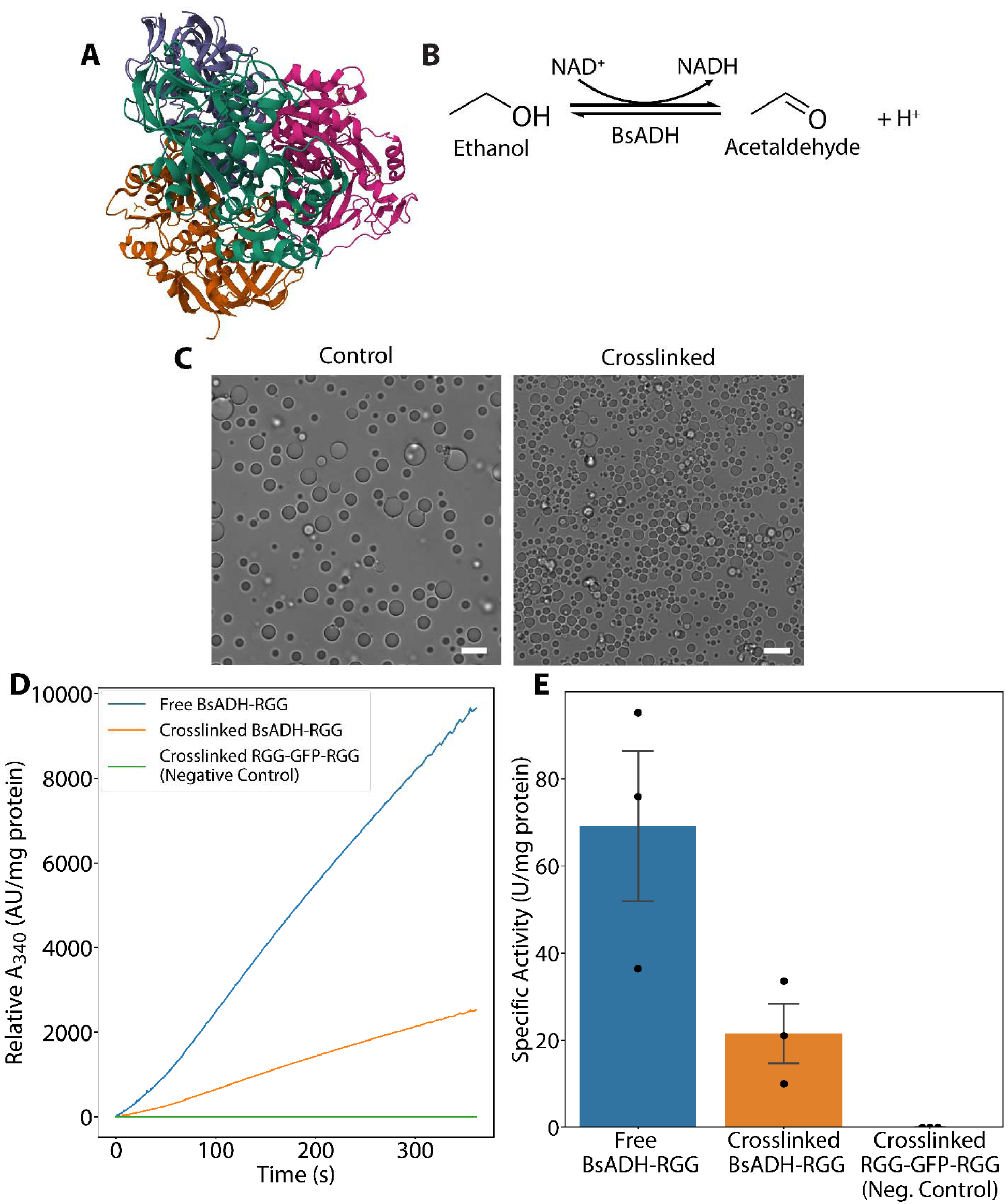
Utilizing crosslinked condensates as a carrier-free method for enzyme immobilization. (A) Protein structure of the homotetrameric BsADH complex determined by x-ray crystallography (PDB: 1RJW). (B) Redox reaction catalyzed by BsADH. Ethanol and other alcohols are oxidized to their respective aldehydes while an NAD^+^ cofactor is reduced to NADH. (C) Representative DIC microscopy image of control (left) and crosslinked (right) BsADH-RGG condensates in a solution containing 50 mM sodium phosphate, pH 8.0, 10 mM 2-mercaptoethanol, 5% (w/v) PEG-8K. Scale bars = 10 µm. (D) Representative BsADH activity measurement by spectrophotometry. The change in A_340_ due to formation of NADH is used to calculate BsADH activity. (E) Bar plots showing measured enzyme activity in free BsADH-RGG (untreated control), crosslinked BsADH-RGG condensates, and RGG-GFP-RGG (secondary negative control). Error bars represent SEM. N = 3 separate batches.

We found that crosslinked BsADH retained 31.1% of its activity compared to free BsADH-RGG (Fig. 7E). We hypothesize that the observed activity loss stemmed from restricted conformational flexibility in the crosslinked enzyme. This hypothesis is supported by an amide hydrogen-deuterium exchange study suggesting that NAD^+^-binding to BsADH subunits induced conformational changes in the enzyme.^52^ Lysines located in the cofactor binding domain of BsADH are susceptible to crosslinking, which could diminish the enzyme’s NAD^+^ binding. Further supporting this hypothesis, crosslinked BsADH-RGG condensates displayed reduced partitioning of fluorescein-conjugated NAD^+^ compared to untreated controls (Supp. Fig. 8). We attempted to rescue some of this activity loss by adding NAD^+^ in 10-fold molar excess to the BsADH-RGG solution before crosslinking, but this did not lead to any appreciable differences (Supp. Fig. 9). Nevertheless, our data demonstrate that we successfully crosslinked BsADH-RGG condensates with partially retained alcohol dehydrogenase activity, confirming at least partial preservation of functional tertiary structure post-crosslinking.

As an alternative, indirect method of immobilization, we also cloned a SpyTagged alcohol dehydrogenase (BsADH-ST3) that could be captured into RGG-SpyCatcher-RGG POMPOMS. However, the fused SpyTag compromised the enzyme’s thermostability and decreased its overall activity (Supp. Fig. 11C), even in the absence of POMPOMS. The enzyme could be captured into POMPOMS (Supp. Fig. 10) but only showed a small amount of residual activity (Supp. Fig. 11A,B). Enzyme immobilization into POMPOMS via SpyTag/SpyCatcher may be viable, but further testing will be needed, perhaps using different enzymes. In all, our results validate POMPOMS as a platform for enzyme immobilization, with future opportunities to improve enzyme activity with further optimization.

### Conclusion

In this work, we established a robust strategy for fabricating protein microparticles by crosslinking biomolecular condensates, overcoming limitations of other techniques to form functional, micrometer-scale, protein-based materials. With POMPOMS, we demonstrated specific applications, including high-affinity molecular capture and enzyme immobilization, validating the platform’s utility and the preservation of functional protein tertiary structure. We also successfully generated core-shell POMPOMS, highlighting the potential for structured material design that could enable a wide array of functionalities.

The process of immobilizing enzymes through POMPOMS formation is related to other carrier-free methods of immobilization, such as crosslinked enzyme aggregates (CLEAs), but our approach offers several benefits.^53,54^ Whereas CLEAs require the use of organic solvents or harsh buffer conditions to aggregate enzymes, POMPOMS take advantage of IDPs to drive phase separation under mild, aqueous conditions.^55^ Unlike CLEAs, POMPOMS form porous and generally spherical materials without requiring the addition of co-aggregants such as BSA or starch, which could help alleviate diffusional limitations and facilitate process design for reactors of different types.^56–58^ Indeed, we demonstrated that POMPOMS retained the permeability characteristic of liquid biomolecular condensates. Furthermore, we demonstrated POMPOMS size control using simple parameters (protein concentration and coalescence time before crosslinking); more advanced techniques such as microfluidics, the addition of macromolecular crowders, or hierarchical emulsion systems could be applied for even finer size control.

By using genetic engineering to integrate the self-assembly behavior of intrinsically disordered proteins with functional folded domains, coupled with standard bioconjugate chemistry, POMPOMS expand the repertoire of protein-based materials for industrial applications. Future work could explore multiplexed functionalities and scalability, positioning POMPOMS as a transformative platform in biological-based materials and biotechnology.

## ASSOCIATED CONTENT

**Supporting Information:** (1) sequences of all proteins used, (2) mathematical derivation of the BsADH specific activity equation, (3) additional experimental details and methods for BsADH activity determination, (4) a representative SDS-PAGE image used to calculate crosslinking yields of RGG-GFP-RGG POMPOMS synthesized using a 5-fold to 200-fold molar excess BS^3^, (5) representative confocal microscopy images showing photobleached RGG-GFP-RGG condensates/POMPOMS, (6) experiments elucidating the kinetics of the BS^3^ crosslinking reaction with RGG-GFP-RGG condensates, (7) representative microscopy images of irregularly-shaped POMPOMS formed as a result of fast reaction kinetics, (8) a comparison of molecular permeability between RGG-GFP-RGG condensates and POMPOMS, (9) microscopy images showing the partitioning of 0.10 and 0.50 µm PS particles into RGG-GFP-RGG condensates, (10) a comparison of binding capacity in RGG-SpyCatcher-RGG POMPOMS batches that were crosslinked with 5-fold to 200-fold molar excess BS^3^, (11) a comparison of fluorescein-NAD^+^ partitioning into BsADH-RGG condensates and BsADH-RGG POMPOMS, (12) comparison of BsADH activity between uncrosslinked BsADH-RGG, crosslinked BsADH-RGG, BsADH-RGG that was crosslinked in the presence of 10-fold molar excess NAD^+^, and crosslinked RGG-GFP-RGG, (13) attempts to immobilize BsADH by capturing them into RGG-SpyCatcher-RGG POMPOMS (PDF).

## Author Contributions

The manuscript was written through contributions of all authors. All authors have given approval to the final version of the manuscript.

## Funding Sources

NJHF grant PC 105-21, NSF grant DMR-2238914.

## Notes

The authors declare no competing financial interests.

## ACKNOWLEDGMENT

This work was supported by NJHF grant PC 105-21 and NSF grant DMR-2238914. A.P. was supported by NIH training grant T32GM135141.

## ABBREVIATIONS

ANOVA: analysis of variance
AU: arbitrary units
BCA: bicinchoninic acid
BS^3^: bis(sulfosuccinimidyl) suberate
BsADH: alcohol dehydrogenase from Geobacillus stearothermophilus
CLEA: crosslinked enzyme aggregate.
CV: column volume
DNA: deoxyribonucleoic acid
EDTA: ethylenediaminetetraacetic acid
FPLC: fast protein liquid chromatography
FRAP: fluorescence recovery after photobleaching
GFP: green fluorescent protein
IDP: intrinsically disordered protein
IPTG: iso-propyl-ß-D-1-thiogalactopyranoside
kDa: kilodalton
LB: lysogeny broth
LLPS: liquid-liquid phase separation
MBP: maltose-binding protein
MES: 2-morpholinoethanesulfonic acid
MW: molecular weight
MWCO: molecular weight cutoff
NA: numerical aperture
NAD+: nicotinamide adenine dinucleotide (oxidized)
NADH: nicotinamide adenine dinucleotide (reduced)
NIR: near infrared
OD: optical density
PEG: polyethylene glycol
POMPOMS: protein-based, self-organized micro-particles of multifunctional significance
PS: polystyrene
*R_h_*: hydrodynamic radius
SDS: sodium dodecyl sulfate
SDS-PAGE: sodium dodecyl sulfate-polyacrylamide gel electrophoresis
SEM: standard error of means
TB: Terrific Broth
T_m_’: apparent melting temperature
TRITC: tetramethylrhodamine isothiocyanate
UCST: upper critical solution temperature
UV: ultra-violet

## Notes

### Competing Interest Statement

The authors have declared no competing interest.

### Summary of Updates

New experiments exploring the effects of crosslinker concentration on microparticle properties. New figure 3 added. Supplemental files added.

