## Supporting Information for "POMPOMS: Crosslinked biomolecular condensates as a versatile platform for multifunctional protein microparticles"

Supplementary Notes

Supplementary Figures 1 to 10

**Supplementary Notes**

Sequences of proteins used in this paper

**RGG-GFP-RGG (RGG-EGFP-RGG) Sequence**

MESNQSNNGGSGNAALNRGGRYVPPHLRGGDGGAAAAASAGGDDRRGGAGGGGYRRGGGNSGGGGGGGYDRGYNDNRDDRDNRGGSGGYGRDRNYEDRGYNGGGGGGGNRGYNNNRGGGGGGYNRQDRGDGGSSNFSRGGYNNRDEGSDNRGSGRSYNNDRRDNGGDGEFLVPRGSMVSKGEELFTGVVPILVELDGDVNGHKFSVSGEGEGDATYGKLTLKFICTTGKLPVPWPTLVTTLTYGVQCFSRYPDHMKQHDFFKSAMPEGYVQERTIFFKDDGNYKTRAEVKFEGDTLVNRIELKGIDFKEDGNILGHKLEYNYNSHNVYIMADKQKNGIKVNFKIRHNIEDGSVQLADHYQQNTPIGDGPVLLPDNHYLSTQSALSKDPNEKRDHMVLLEFVTAAGITLGMDELYKEFLVPRGSMESNQSNNGGSGNAALNRGGRYVPPHLRGGDGGAAAAASAGGDDRRGGAGGGGYRRGGGNSGGGGGGGYDRGYNDNRDDRDNRGGSGGYGRDRNYEDRGYNGGGGGGGNRGYNNNRGGGGGGYNRQDRGDGGSSNFSRGGYNNRDEGSDNRGSGRSYNNDRRDNGGDGLEHHHHHH*

**RGG-SpyCatcher-RGG (RGG-SpyCatcher003-RGG) Sequence**

MESNQSNNGGSGNAALNRGGRYVPPHLRGGDGGAAAAASAGGDDRRGGAGGGGYRRGGGNSGGGGGGGYDRGYNDNRDDRDNRGGSGGYGRDRNYEDRGYNGGGGGGGNRGYNNNRGGGGGGYNRQDRGDGGSSNFSRGGYNNRDEGSDNRGSGRSYNNDRRDNGGDGGSGESGSGVTTLSGLSGEQGPSGDMTTEEDSATHIKFSKRDEDGRELAGATMELRDSSGKTISTWISDGHVKDFYLYPGKYTFVETAAPDGYEVATPIEFTVNEDGQVTVDGEATEGDAHTGSGESGSGESNQSNNGGSGNAALNRGGRYVPPHLRGGDGGAAAAASAGGDDRRGGAGGGGYRRGGGNSGGGGGGGYDRGYNDNRDDRDNRGGSGGYGRDRNYEDRGYNGGGGGGGNRGYNNNRGGGGGGYNRQDRGDGGSSNFSRGGYNNRDEGSDNRGSGRSYNNDRRDNGGDGLEHHHHHH*

**SZ2-GFP (SZ2-GGGGS-TEV site-mEGFP) Sequence**

MARNAYLRKKIARLKKDNLQLERDEQNLEKIIANLRDEIARLENEVASHEQGGGGSENLYFQVSKGEELFTGVVPILVELDGDVNGHKFSVSGEGEGDATYGKLTLKFICTTGKLPVPWPTLVTTLTYGVQCFSRYPDHMKQHDFFKSAMPEGYVQERTIFFKDDGNYKTRAEVKFEGDTLVNRIELKGIDFKEDGNILGHKLEYNYNSHNVYIMADKQKNGIKVNFKIRHNIEDGSVQLADHYQQNTPIGDGPVLLPDNHYLSTQSKLSKDPNEKRDHMVLLEFVTAAGITLGMDELYKHHHHHH*

**GFP-SpyTag (6His-mEGFP-SpyTag003) Sequence**

MHHHHHHVSKGEELFTGVVPILVELDGDVNGHKFSVSGEGEGDATYGKLTLKFICTTGKLPVPWPTLVTTLTYGVQCFSRYPDHMKQHDFFKSAMPEGYVQERTIFFKDDGNYKTRAEVKFEGDTLVNRIELKGIDFKEDGNILGHKLEYNYNSHNVYIMADKQKNGIKVNFKIRHNIEDGSVQLADHYQQNTPIGDGPVLLPDNHYLSTQSKLSKDPNEKRDHMVLLEFVTAAGITLGMDELYKGGSGRGVPHIVMVDAYKRYK *

**SpyTag-GFP (SpyTag003-mEGFP-6His) Sequence**

MRGVPHIVMVDAYKRYKGSGESGVSKGEELFTGVVPILVELDGDVNGHKFSVSGEGEGDATYGKLTLKFICTTGKLPVPWPTLVTTLTYGVQCFSRYPDHMKQHDFFKSAMPEGYVQERTIFFKDDGNYKTRAEVKFEGDTLVNRIELKGIDFKEDGNILGHKLEYNYNSHNVYIMADKQKNGIKVNFKIRHNIEDGSVQLADHYQQNTPIGDGPVLLPDNHYLSTQSKLSKDPNEKRDHMVLLEFVTAAGITLGMDELYKLEHHHHHH*

**RGG-RGG Sequence**

MESNQSNNGGSGNAALNRGGRYVPPHLRGGDGGAAAAASAGGDDRRGGAGGGGYRRGGGNSGGGGGGGYDRGYNDNRDDRDNRGGSGGYGRDRNYEDRGYNGGGGGGGNRGYNNNRGGGGGGYNRQDRGDGGSSNFSRGGYNNRDEGSDNRGSGRSYNNDRRDNGGDGEFGKLMESNQSNNGGSGNAALNRGGRYVPPHLRGGDGGAAAAASAGGDDRRGGAGGGGYRRGGGNSGGGGGGGYDRGYNDNRDDRDNRGGSGGYGRDRNYEDRGYNGGGGGGGNRGYNNNRGGGGGGYNRQDRGDGGSSNFSRGGYNNRDEGSDNRGSGRSYNNDRRDNGGDGLEHHHHHH*

**MBP-GFP-RGG Sequence**

MKIEEGKLVIWINGDKGYNGLAEVGKKFEKDTGIKVTVEHPDKLEEKFPQVAATGDGPDIIFWAHDRFGGYAQSGLLAEITPDKAFQDKLYPFTWDAVRYNGKLIAYPIAVEALSLIYNKDLLPNPPKTWEEIPALDKELKAKGKSALMFNLQEPYFTWPLIAADGGYAFKYENGKYDIKDVGVDNAGAKAGLTFLVDLIKNKHMNADTDYSIAEAAFNKGETAMTINGPWAWSNIDTSKVNYGVTVLPTFKGQPSKPFVGVLSAGINAASPNKELAKEFLENYLLTDEGLEAVNKDKPLGAVALKSYEEELVKDPRIAATMENAQKGEIMPNIPQMSAFWYAVRTAVINAASGRQTVDEALKDAQTNSSSNNNNNNNNNNLGETVRFQSMVSKGEELFTGVVPILVELDGDVNGHKFSVSGEGEGDATYGKLTLKFICTTGKLPVPWPTLVTTLTYGVQCFSRYPDHMKQHDFFKSAMPEGYVQERTIFFKDDGNYKTRAEVKFEGDTLVNRIELKGIDFKEDGNILGHKLEYNYNSHNVYIMADKQKNGIKVNFKIRHNIEDGSVQLADHYQQNTPIGDGPVLLPDNHYLSTQSKLSKDPNEKRDHMVLLEFVTAAGITLGMDELYKGGGSENLYFQGEFGKLMESNQSNNGGSGNAALNRGGRYVPPHLRGGDGGAAAAASAGGDDRRGGAGGGGYRRGGGNSGGGGGGGYDRGYNDNRDDRDNRGGSGGYGRDRNYEDRGYNGGGGGGGNRGYNNNRGGGGGGYNRQDRGDGGSSNFSRGGYNNRDEGSDNRGSGRSYNNDRRDNGGDGLEHHHHHH*

**BsADH-RGG Sequence**

MKAAVVEQFKEPLKIKEVEKPTISYGEVLVRIKACGVCHTDLHAAHGDWPVKPKLPLIPGHEGVGIVEEVGPGVTHLKVGDRVGIPWLYSACGHCDYCLSGQETLCEHQKNAGYSVDGGYAEYCRAAADYVVKIPDNLSFEEAAPIFCAGVTTYKALKVTGAKPGEWVAIYGIGGLGHVAVQYAKAMGLNVVAVDIGDEKLELAKELGADLVVNPLKEDAAKFMKEKVGGVHAAVVTAVSKPAFQSAYNSIRRGGACVLVGLPPEEMPIPIFDTVLNGIKIIGSIVGTRKDLQEALQFAAEGKVKTIIEVQPLEKINEVFDRMLKGQINGRVVLTLEDKGMESNQSNNGGSGNAALNRGGRYVPPHLRGGDGGAAAAASAGGDDRRGGAGGGGYRRGGGNSGGGGGGGYDRGYNDNRDDRDNRGGSGGYGRDRNYEDRGYNGGGGGGGNRGYNNNRGGGGGGYNRQDRGDGGSSNFSRGGYNNRDEGSDNRGSGRSYNNDRRDNGGDGLEHHHHHH*

**BsADH Sequence**

MKAAVVEQFKEPLKIKEVEKPTISYGEVLVRIKACGVCHTDLHAAHGDWPVKPKLPLIPGHEGVGIVEEVGPGVTHLKVGDRVGIPWLYSACGHCDYCLSGQETLCEHQKNAGYSVDGGYAEYCRAAADYVVKIPDNLSFEEAAPIFCAGVTTYKALKVTGAKPGEWVAIYGIGGLGHVAVQYAKAMGLNVVAVDIGDEKLELAKELGADLVVNPLKEDAAKFMKEKVGGVHAAVVTAVSKPAFQSAYNSIRRGGACVLVGLPPEEMPIPIFDTVLNGIKIIGSIVGTRKDLQEALQFAAEGKVKTIIEVQPLEKINEVFDRMLKGQINGRVVLTLEDKLEHHHHHH*

**BsADH-SpyTag Sequence**

MKAAVVEQFKEPLKIKEVEKPTISYGEVLVRIKACGVCHTDLHAAHGDWPVKPKLPLIPGHEGVGIVEEVGPGVTHLKVGDRVGIPWLYSACGHCDYCLSGQETLCEHQKNAGYSVDGGYAEYCRAAADYVVKIPDNLSFEEAAPIFCAGVTTYKALKVTGAKPGEWVAIYGIGGLGHVAVQYAKAMGLNVVAVDIGDEKLELAKELGADLVVNPLKEDAAKFMKEKVGGVHAAVVTAVSKPAFQSAYNSIRRGGACVLVGLPPEEMPIPIFDTVLNGIKIIGSIVGTRKDLQEALQFAAEGKVKTIIEVQPLEKINEVFDRMLKGQINGRVVLTLEDKGGSGRGVPHIVMVDAYKRYKLEHHHHHH*

**Supplementary Note 1: Derivation of the BsADH specific activity equation**

The following reaction was used to measure the specific activity of BsADH by continuous spectrophotometry at 340 nm:

$$\text{Ethanol}+\text{NAD}^{+}\underset{\to}{\text{BsADH}}\text{Acetaldehyde}+\text{NADH}$$

One unit of BsADH activity was defined as the conversion of 1 µmol NAD^+^ to NADH per minute under the reaction conditions. The BsADH activity is then normalized to the amount of protein added to the reaction cuvette to calculate the specific activity:

$$\text{Specific Activity (Units/}\text{mg}_{\text{protein}}\text{)}=\frac{\text{μmol}_{\text{NADH}} \text{min}^{-1}}{m_{\text{protein, rxn}}}$$

(1)

where m_protein, rxn_ is the mass of protein added to the reaction cuvette.

From the Beer-Lambert Law:

$$A=\epsilon cl$$

(2)

where $A$ is the UV absorbance of NADH at 340 nm,

$\epsilon$ is the molar absorptivity of NADH at 340 nm (6.22 mM^-1^ cm^-1^),

$c$ is the concentration of NADH in solution,

and $l$ is the path length of the cuvette (1 cm).

The change in NADH concentrated can be equated to the change in absorbance. Therefore, the Beer-Lambert Law can be rewritten as:

$$\Delta c=\frac{\Delta A}{\epsilon l}$$

(3)

where $\Delta c$ is the change in NADH concentration (mM min^-1^),

and $\Delta A$ is the change in absorbance (min^-1^).

Eq. (3) can be multiplied by the total reaction volume to find change in amount of NADH, which can be substituted into eq. (1):

$$\text{Specific Activity (Units/}\text{mg}_{\text{protein}}\text{)}=\frac{\Delta c*V_{\text{assay}}}{m_{\text{protein, rxn}}}=\frac{\Delta A*V_{\text{assay}}}{\epsilon*l*m_{\text{protein, rxn}}}$$

(4)

where V_Assay_ is the total reaction volume in the cuvette (mL).

**Supplementary Note 2: Determination of BsADH activity**

All experiments were performed alongside at least one negative control (buffer only), which allowed for the correction of any changes in any nonspecific changes in A_340_. Eq. (4) can be rewritten as:

$$\text{Specific Activity (Units/}\text{mg}_{\text{protein}}\text{)}=\frac{(\Delta A_{340, \text{Sample}}-\Delta A_{340, \text{Blank}})*V_{\text{Assay}}}{\epsilon*l*m_{\text{protein, rxn}}}$$

(5)

where ΔA_340_ is the average change in A_340_ per minute derived from a linear fit of the raw absorbance data (min^-1^).

Eq. (5) uses the mass of protein added to the reaction cuvette (m_protein, rxn_) to normalize the enzyme activity. However, the mass of protein added to the cuvette is difficult to measure directly. Instead, the protein mass can be calculated from the mass concentration of the protein solution and the volume of protein solution added to the cuvette. Eq. (5) can be rewritten as the following:

$$\text{Specific Activity (Units/}\text{mg}_{\text{protein}}\text{)}=\frac{(\Delta A_{340, \text{Sample}}-\Delta A_{340, \text{Blank}})*V_{\text{Assay}}}{\epsilon*l*V_{\text{protein}}*\rho_{\text{protein}}}$$

(6)

where V_protein_ is the volume of protein solution added to the cuvette (mL),

and ρ_protein_ is the mass concentration of protein added to the cuvette (mg/mL).

In Eq. (6), V_Assay_, ϵ, l, V_protein_, and ρ_protein_ are known values, leaving only the ΔA_340_ values to be determined by the enzyme assay.

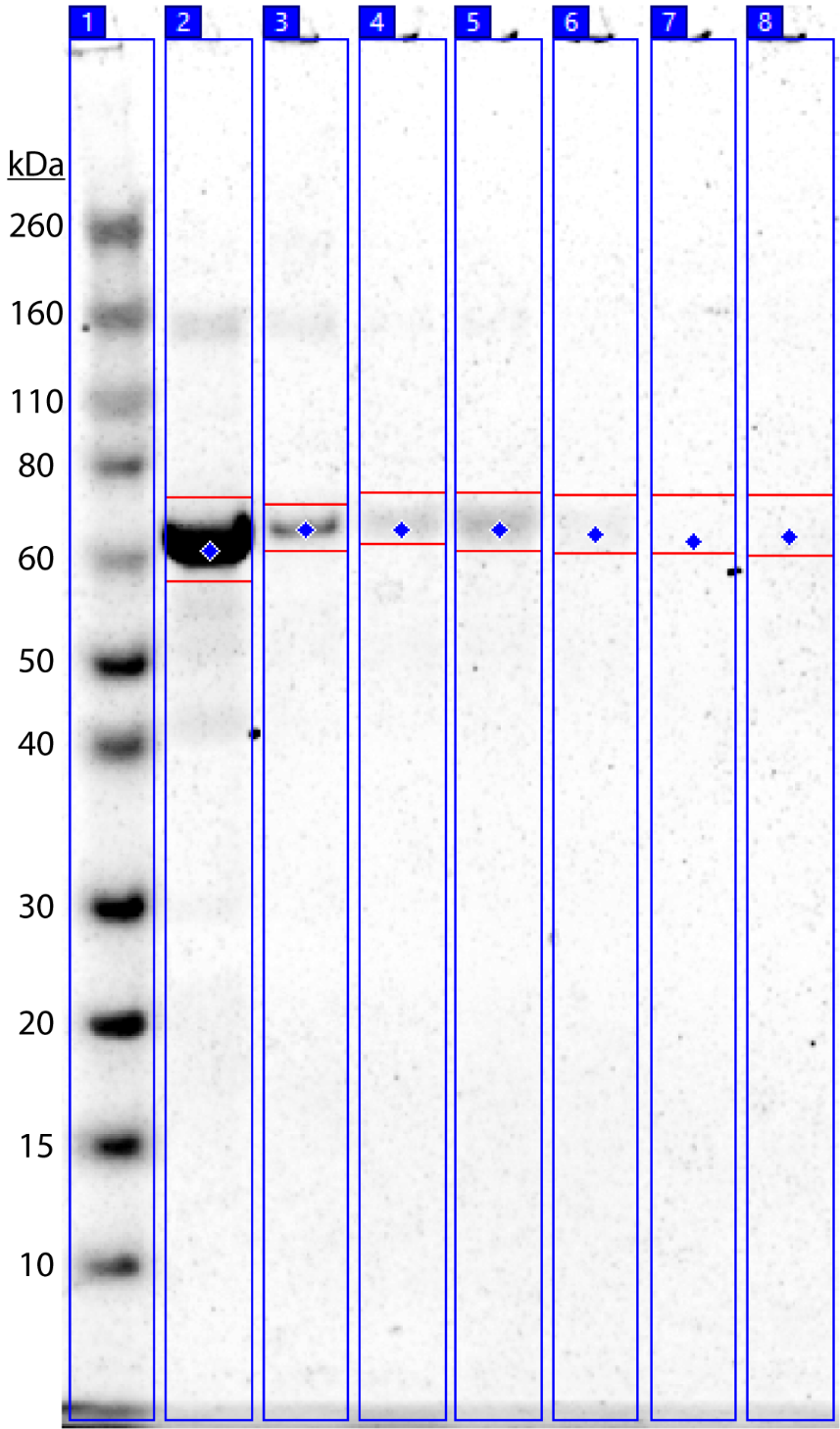

**Supplementary Figure 1. Representative near-infrared (NIR) fluorescence image of Coomassie-stained SDS-PAGE gel used to calculate crosslinking yields of RGG-GFP-RGG POMPOMS batches prepared with 0-200X molar excess BS^3^.** The gel shows, from left to right: (1) protein standard ladder, (2) 0X (uncrosslinked) RGG-GFP-RGG, (3-8) RGG-GFP-RGG crosslinked with (3) 5X, (4) 10X, (5) 20X, (6) 50X, (7) 100X, (8) 200X molar excess BS^3^. The blue diamond shows the center of the initial selected area, and the horizontal red lines represent the edges of the analyzed area after manual adjustment. Crosslinking yield was calculated by comparing the monomer bands corresponding to RGG-GFP-RGG (MW: 63.3 kDa) from crosslinked samples (lanes 3-8) to an uncrosslinked control (lane 2).

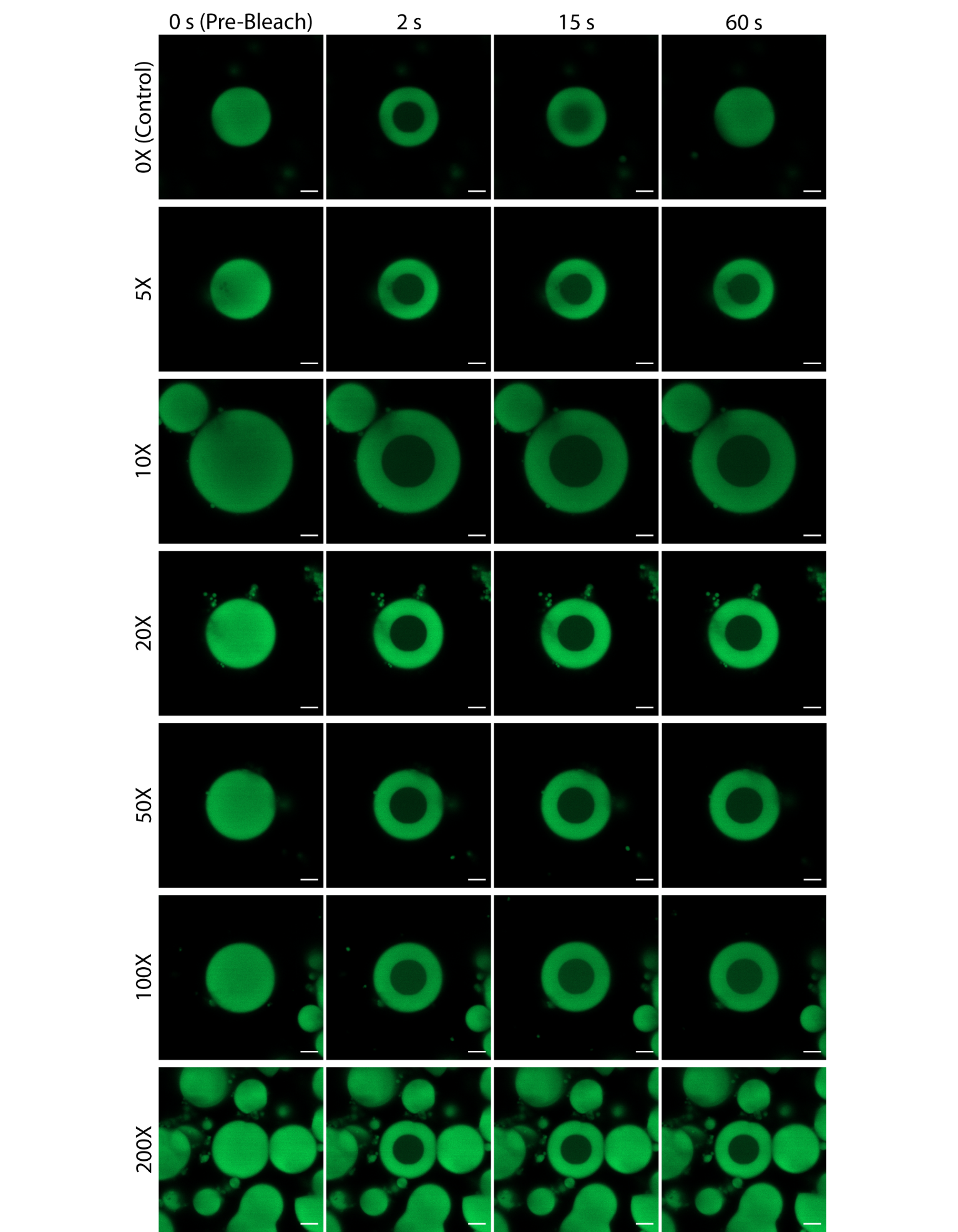

**Supplementary Figure 2. Representative confocal microscopy images from fluorescence recovery after photobleaching (FRAP) experiments comparing RGG-GFP-RGG condensates (control) to RGG-GFP-RGG POMPOMS crosslinked with 5-200X molar excess BS^3^.** Even at the lowest tested concentration of BS^3^ (5X), RGG-GFP-RGG POMPOMS show limited fluorescence recovery after photobleaching. In addition, the boundaries of the photobleached area remain distinct for each crosslinked condition. On the other hand, the photobleached area in uncrosslinked RGG-GFP-RGG (0X) recovers its fluorescence, and the photobleached area’s boundaries dissipate because of dynamic exchange. Scale bar = 10 µm.

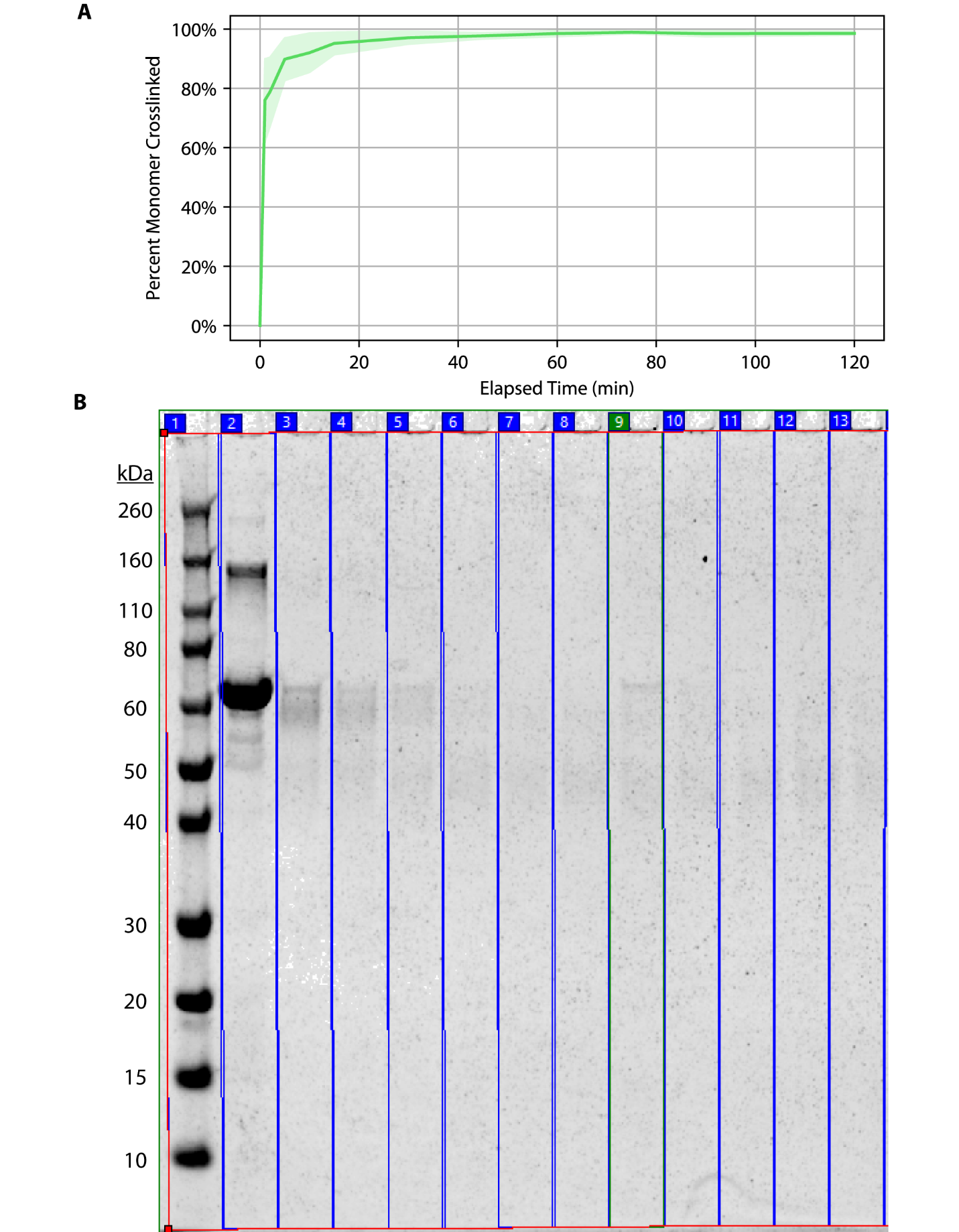

**Supplementary Figure 3. Kinetics of BS^3^ crosslinking reaction.** (A) Summary results of three trials. The banded area represents the standard error of means for each measured timepoint. (B) Representative NIR fluorescence image of the SDS-PAGE gel used to calculate the amount of RGG-GFP-RGG monomer crosslinked at each timepoint by comparing to the band intensity at time 0 (uncrosslinked control). The gel shows, from left to right: (1) protein standard ladder, (2) uncrosslinked RGG-GFP-RGG control, (3-13) RGG-GFP-RGG samples after incubating with 20X molar excess BS^3^ for (3) 1 min, (4) 2 min, (5) 5 min, (6) 10 min, (7) 15 min, (8) 30 min (9) 45 min, (10) 60 min, (11) 75 min, (12) 90 min, (13) 120 min.

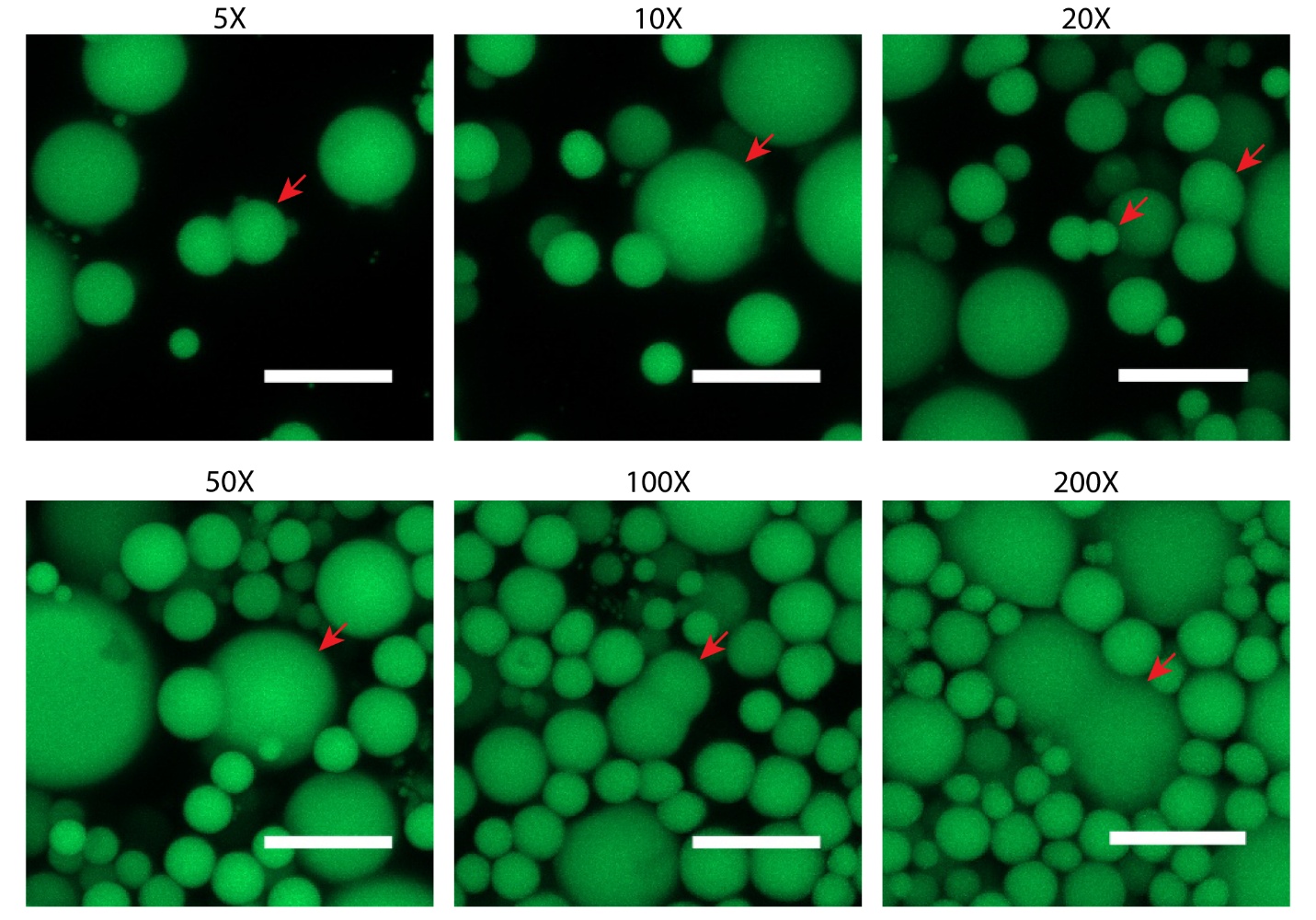

**Supplementary Figure 4. Representative microscopy images of irregularly shaped POMPOMS from batches crosslinked with 5X-200X BS^3^.** Irregularly shaped POMPOMS are indicated by the red arrows. Scale bars = 10 µm.

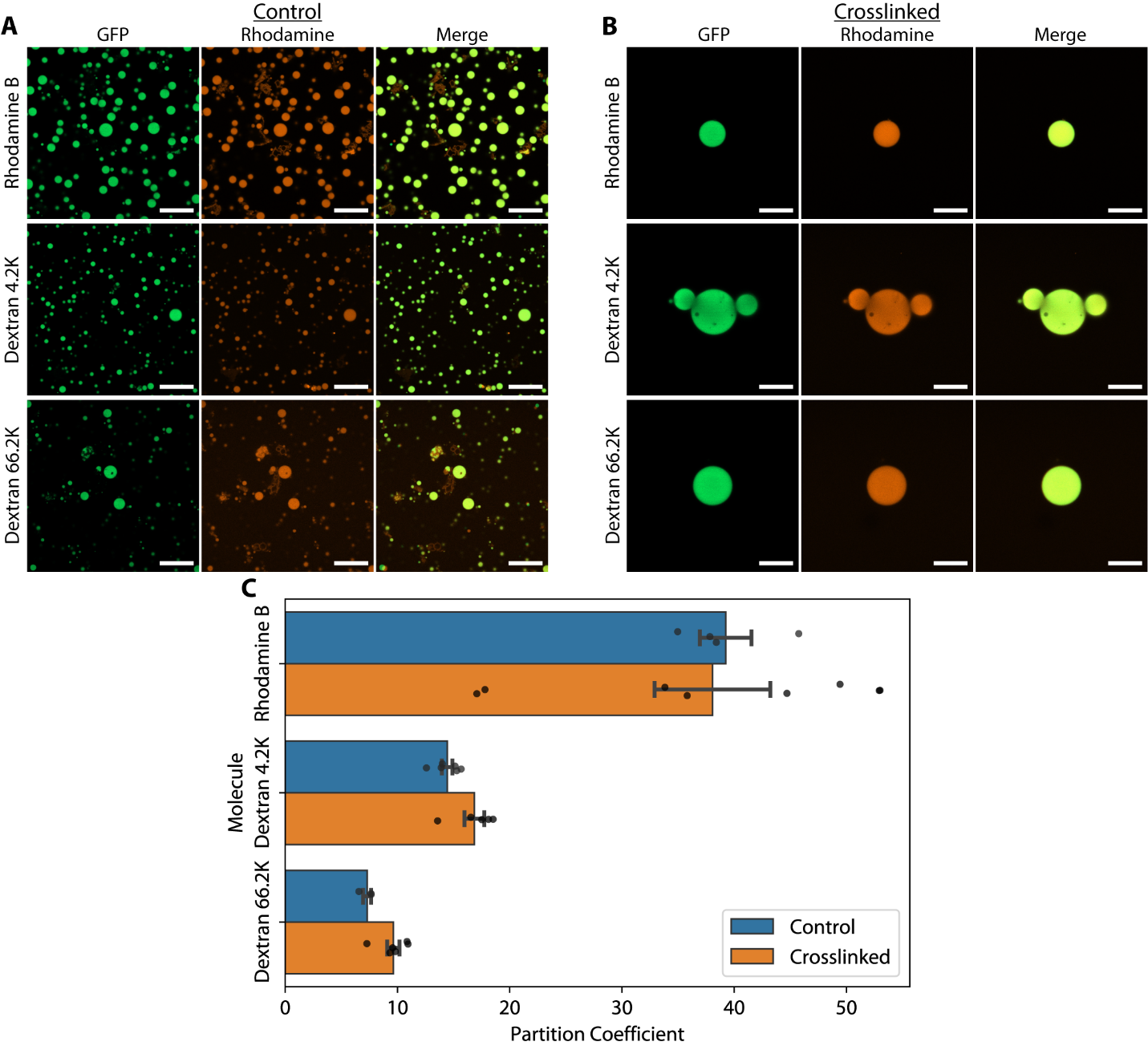

**Supplementary Figure 5.** **RGG-GFP-RGG POMPOMS maintain the molecular permeability of uncrosslinked RGG-GFP-RGG condensates.** (A) Representative microscopy images demonstrating colocalization of rhodamine B and TRITC-labeled dextrans inside liquid RGG-GFP-RGG condensates. (B) Representative microscopy images demonstrating colocalization of rhodamine B and TRITC-labeled dextrans inside crosslinked RGG-GFP-RGG microparticles. These images are the same that appear in Figure 4A of the main text. (C) Bar plot comparing rhodamine and TRITC-labeled dextran partition coefficients between control (untreated) and crosslinked RGG-GFP-RGG condensates. The differences between partition coefficients for control and crosslinked RGG-GFP-RGG condensates was not found to be significantly different when compared by a one-way ANOVA (p = 0.704, SS = 5201.03, df = 2, F = 42.612). Error bars represent SEM. For all microscopy images, scale bar = 20 µm.

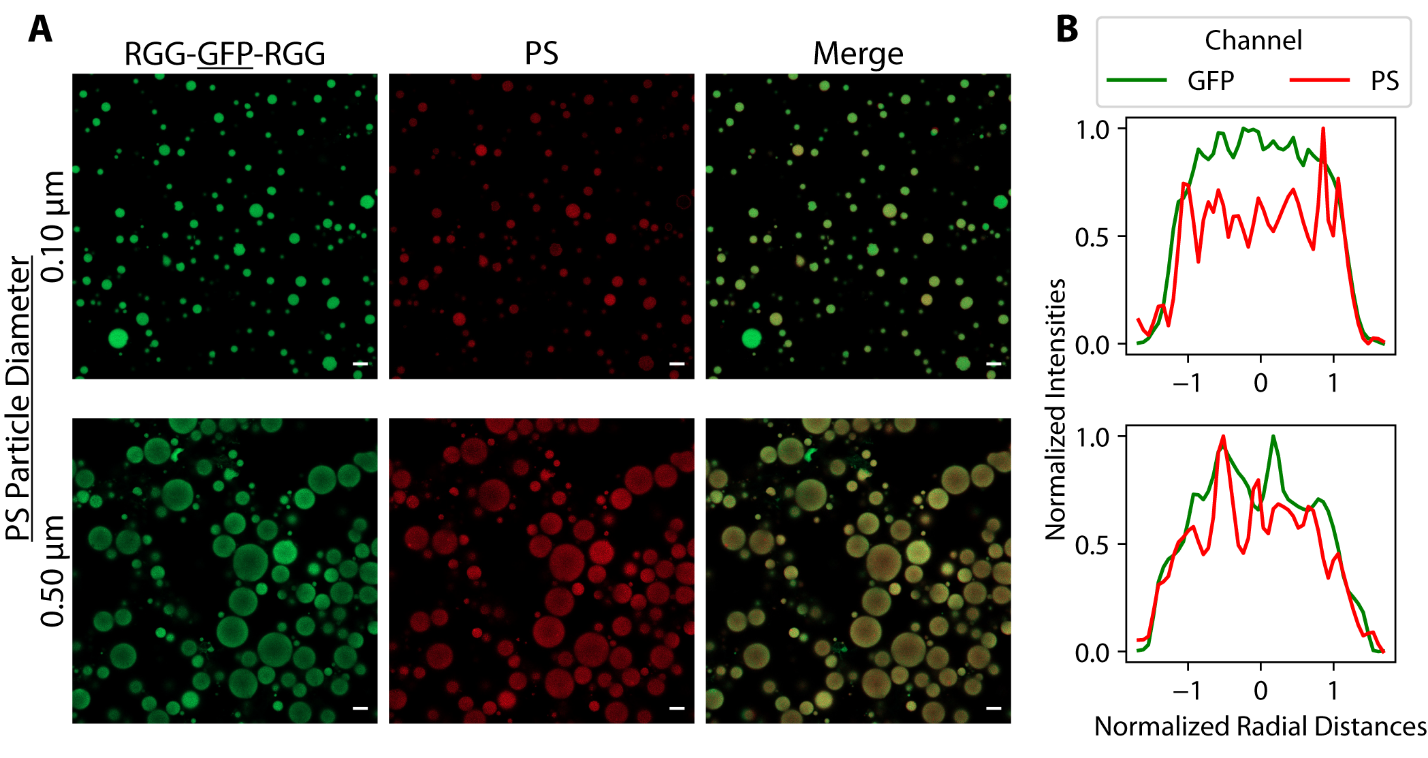

**Supplementary Figure 6. Partitioning of 0.10 and 0.50 µm PS particles into uncrosslinked RGG-GFP-RGG condensates.** (A) Representative confocal microscopy images of RGG-GFP-RGG mixed with 0.10 µm diameter (0.01% volume fraction) and 0.50 µm diameter (0.1% volume fraction) carboxyl-modified polystyrene (PS) particles. PS particles were red fluorescent. Scale bars = 5 µm. (B) Line profiles of microscopy images demonstrating colocalization of GFP and PS particles.

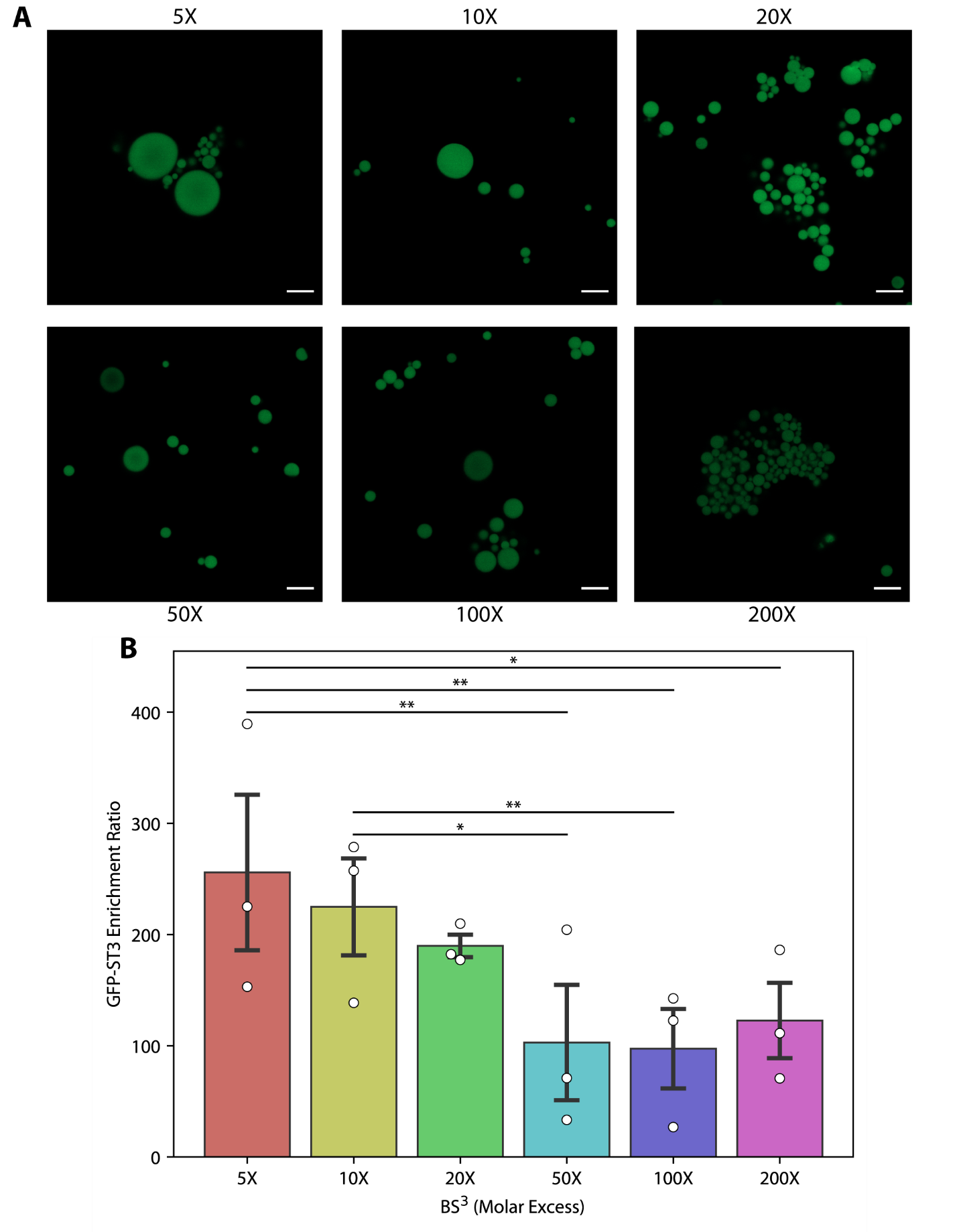

**Supplementary Figure 7. Higher BS^3^ concentrations can reduce the binding capacity of RGG-SpyCatcher-RGG POMPOMS.** (A) Representative confocal microscopy of RGG-Spycatcher-RGG POMPOMS after incubation with GFP-SpyTag (GFP-ST3) followed by several rounds of washing. Scale bars = 10 µm. (B) Bar plot showing quantified results of image analysis. The enrichment ratio (a measure of binding capacity) sharply decreases for POMPOMS prepared using BS^3^ concentrations above 20-fold molar excess. Error bars represent SEM. *** : p < 0.05, **: p < 0.01 when analyzed by a *post-hoc* Tukey test. N = 3 batches.

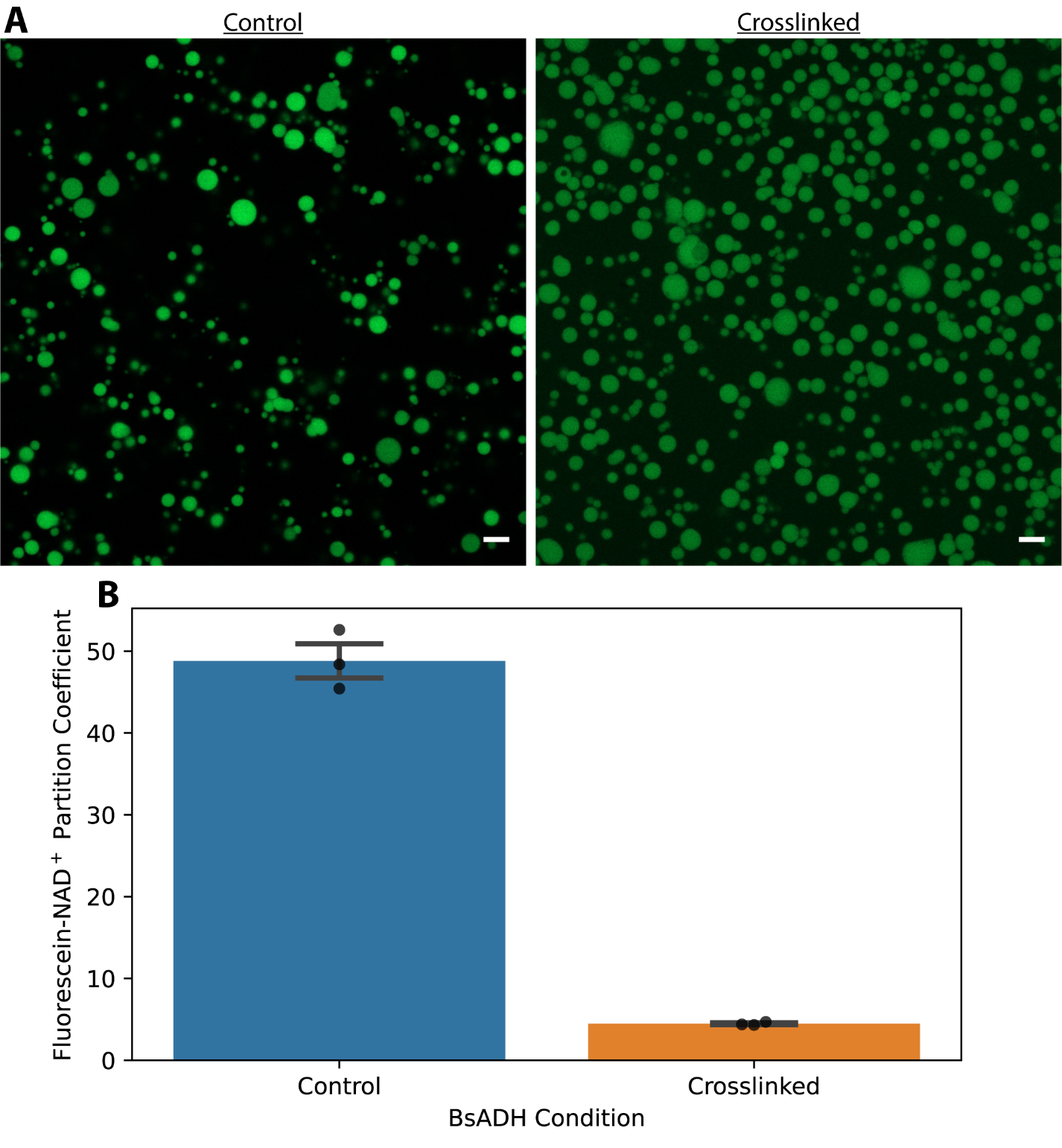

**Supplementary Figure 8. Partitioning of fluorescein-NAD^+^ into control BsADH-RGG condensates and BsADH-RGG POMPOMS.** (A) Representative confocal microscopy images of untreated (left) and crosslinked (right) BsADH-RGG condensates showing varying degrees of fluorescein-NAD^+^ partitioning. Scale bars = 5 µm. (B) Bar plot comparing partition coefficients of fluorescein-NAD+ between control (untreated) and crosslinked BsADH-RGG condensates. Error bars represent SEM. N = 3 images for each condition.

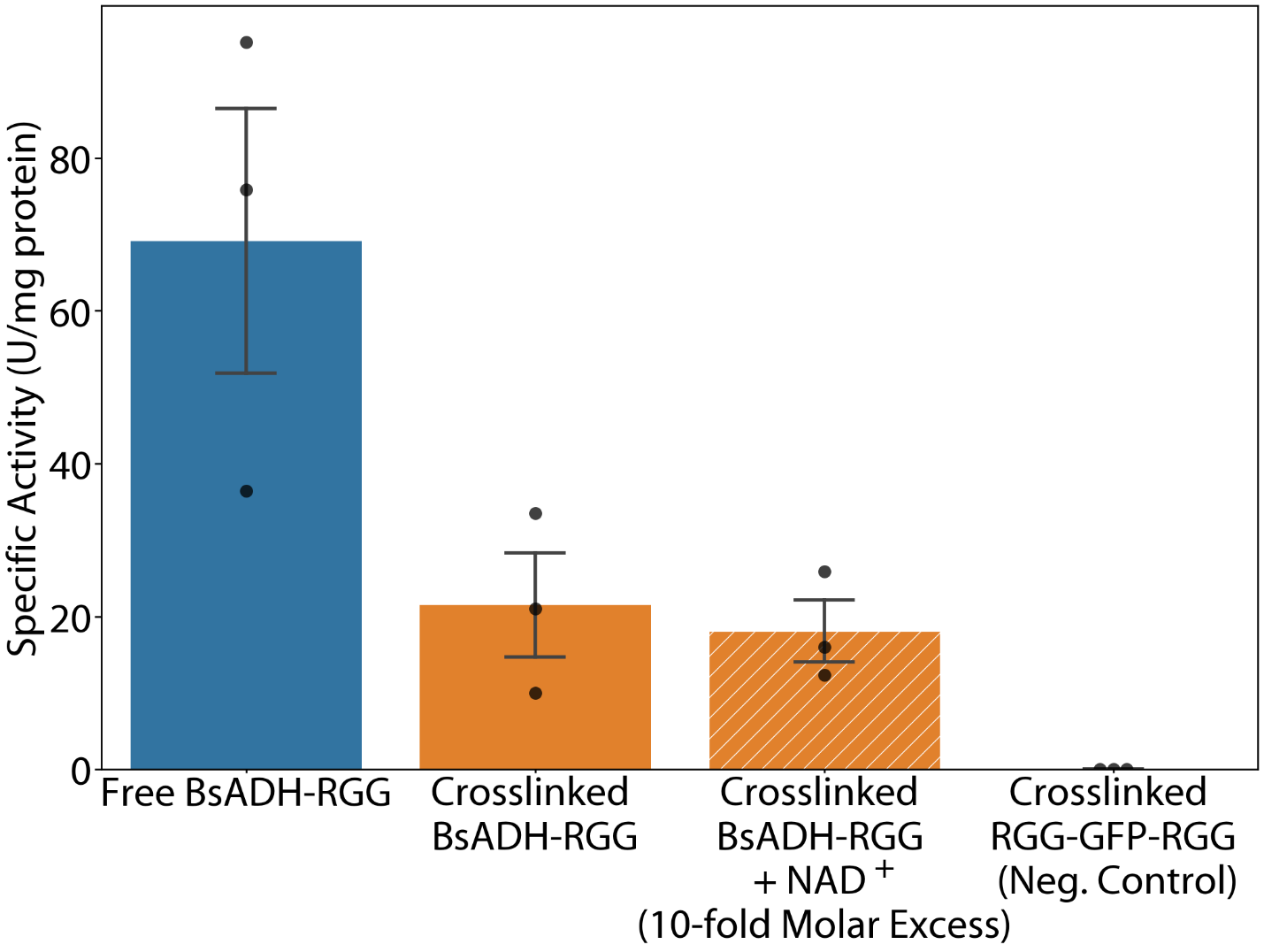

**Supplementary Figure 9. Crosslinking BsADH-RGG in the presence of 10-fold molar excess NAD^+^ did not rescue enzymatic activity.** To attempt to rescue some enzyme activity in crosslinked BsADH-RGG condensates, NAD^+^ was added to the BsADH-RGG solution in 10-fold molar excess before crosslinking. However, this did not lead to any appreciable differences in specific activity after crosslinking. Error bars represent SEM, N = 3 separate batches for each condition. The data for free BsADH-RGG, crosslinked BsADH-RGG, and crosslinked RGG-GFP-RGG are identical to those displayed in Fig. 7E of the main text.

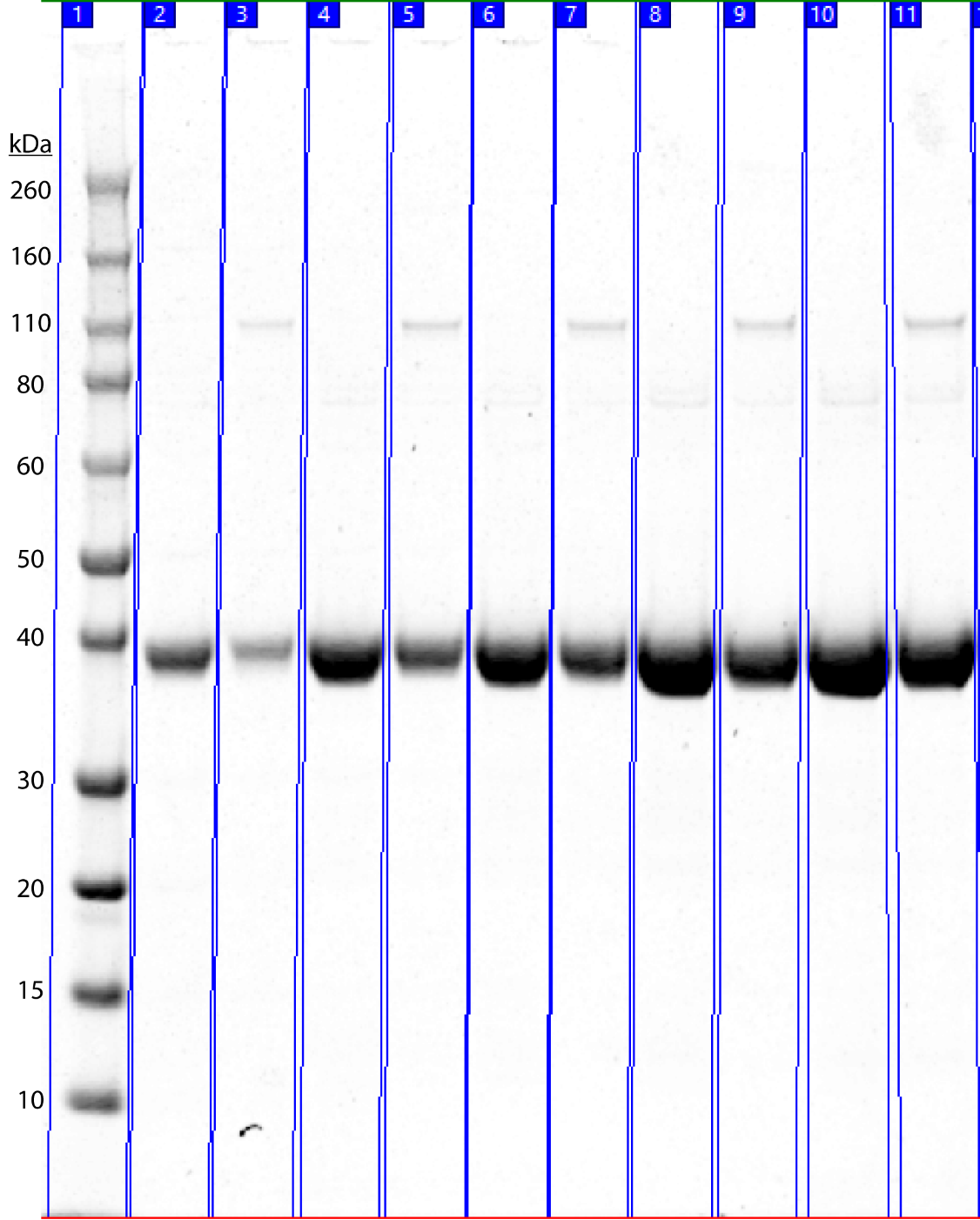

**Supplementary Figure 10. SDS-PAGE gel showing capture of BsADH-SpyTag into RGG-SpyCatcher-RGG POMPOMS.** The gel shows, from left to right: (1) protein standard ladder, (2) 0.2:1 mole ratio BsADH-SpyTag solution (0.16 mg/mL) before conjugation and (3) supernatant after conjugation with RGG-SpyCatcher-RGG POMPOMS, (4) 0.4:1 mole ratio BsADH-SpyTag solution (0.32 mg/mL) before conjugation and (5) after conjugation with RGG-SpyCatcher-RGG POMPOMS, (6) 0.6:1 mole ratio BsADH-SpyTag solution (0.48 mg/mL) before conjugation and (7) after conjugation with RGG-SpyCatcher-RGG POMPOMS, (8) 0.8:1 mole ratio BsADH-SpyTag solution (0.63 mg/mL) before conjugation and (9) after conjugation with RGG-SpyCatcher-RGG POMPOMS, and (10) 1:1 mole ratio BsADH solution (0.79 mg/mL) before conjugation and (11) after conjugation with RGG-SpyCatcher-RGG POMPOMS. BsADH-SpyTag MW: 39.6 kDa.

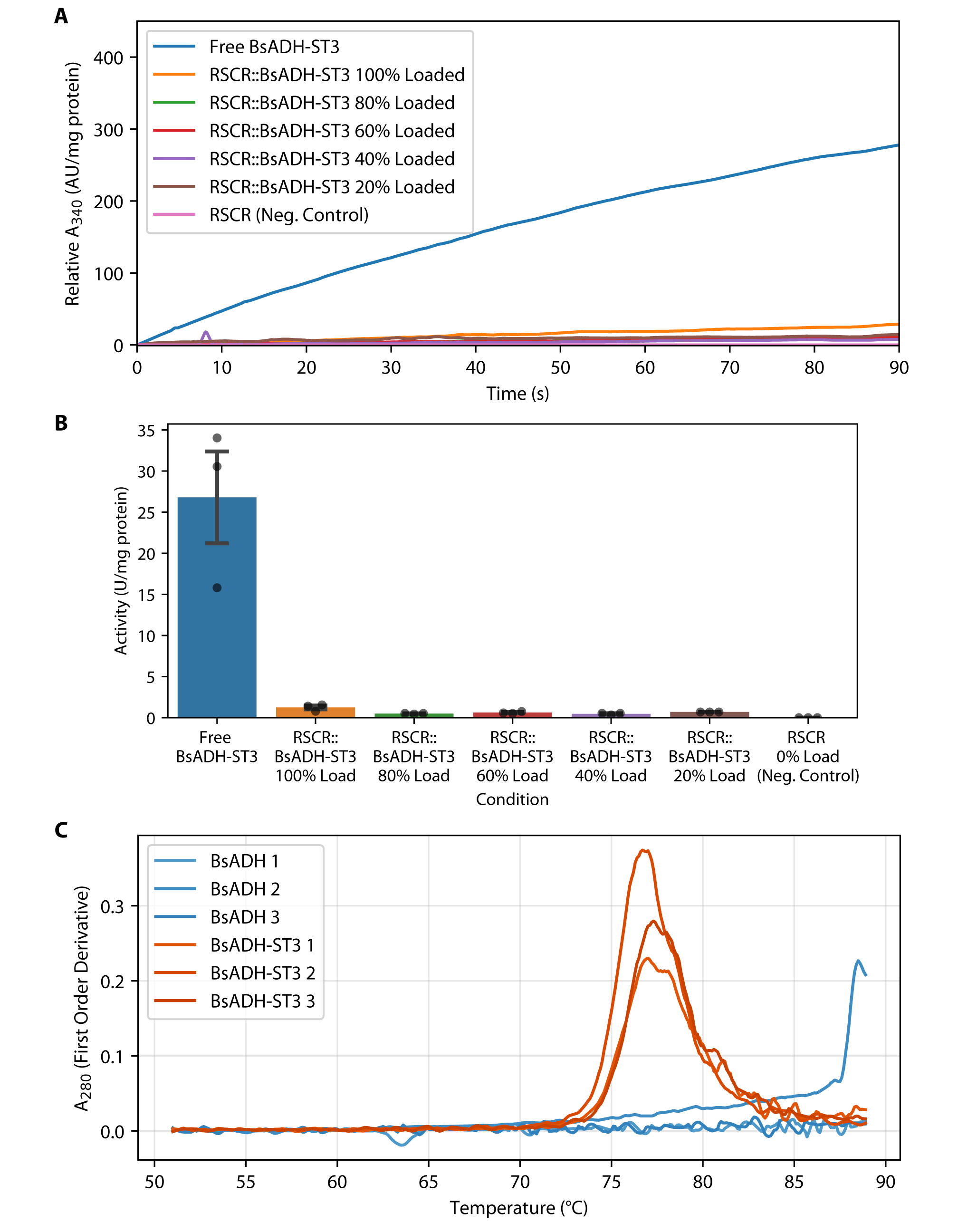

**Supplementary Figure 11. Immobilization of SpyTagged BsADH into POMPOMS.** (A) Representative BsADH-SpyTag activity measurement by spectrophotometry for RGG-SpyCatcher-RGG (RSCR) POMPOMS with various loadings of BsADH-SpyTag (BsADH-ST3). The change in A_340_ due to formation of NADH is used to calculate BsADH-ST3 activity. (B) Bar plots showing the measured activity of BsADH-ST3 in free and immobilized form. RGG-SpyCatcher-RGG (RSCR) POMPOMS that had captured BsADH-ST3 showed some catalytic activity, but it was substantially lower compared to the free enzyme. Error bars represent SEM, and N = 3 separate batches. (C) Comparing thermal stability of wild-type BsADH to BsADH-ST3 by UV-Vis spectroscopy. BsADH/BsADH-SpyTag was heated from 50°C to 90°C and the absorbance at 280 nm was measured. From the data, this first order derivative plot was used to determine apparent melting temperatures (T_m_) using the peak values. BsADH-ST3 samples showed an average T_m_ of 77.0°C, while BsADH did not show clear signs of denaturation at this temperature range. N = 3 replicates for each protein.
